# Photomechanical engineering of immunogenic whole-cell vaccines by laser-induced vapor nanobubbles for bladder cancer prophylaxis

**DOI:** 10.64898/2026.11.06.764866

**Authors:** Núria Sabando-García, Ainhoa G. Caelles, Noemí Traba, Joan Pagès, Cecilia Cabrera, Ana Garcia Hidalgo, Jana Ramon, Deep Punj, Kevin Braeckmans, Annabel F. Valledor, Gertruda Chomiciute, Ibane Abasolo, Samuel Sánchez, Esther Julián, Juan C. Fraire

**Affiliations:** Genetics and Microbiology Department, Faculty of Biosciences, Autonomous University of Barcelona, 08193 Bellaterra, Barcelona, Spain; Institute for Bioengineering of Catalonia (IBEC), Barcelona Institute of Science and Technology (BIST), Baldiri i Reixac 10-12, 08028 Barcelona Spain; Nanobots Therapeutics S.L. Carrer Baldiri i Reixac, 4, 08028 Barcelona, Spain; Department of Biochemistry and Physiology, Faculty of Pharmacy and Food Sciences, University of Barcelona, Av. Diagonal 643, 08028 Barcelona, Spain; Institute for Health Science Research Germans Trias i Pujol (IGTP), Carretera de Canyet s/n, 08916 Badalona, Barcelona, Spain; IrsiCaixa, Hospital Germans Trias i Pujol, Carretera de Canyet s/n, 08916 Badalona, Barcelona, Spain; Laboratory for General Biochemistry and Physical Pharmacy, Faculty of Pharmaceutical Sciences, Ghent University, 9000 Ghent, Belgium; Department of Cell Biology, Physiology and Immunology, Faculty of Biology, University of Barcelona, 08028 Barcelona, Spain; Institute of Biomedicine of the University of Barcelona (IBUB), Av. Diagonal 643, 08028 Barcelona, Spain; Institute for Advanced Chemistry of Catalonia (IQAC-CSIC), Jordi Girona 20, 08034 Barcelona, Spain; Networking Research Center on Bioengineering, Biomaterials and Nanomedicine (CIBER-BBN), Instituto de Salud Carlos III, 08034 Barcelona, Spain; Catalan Institute for Research and Advanced Studies (ICREA), Passeig de Lluís Companys 23, 08010, Barcelona, Spain

**Keywords:** vapor nanobubbles, photomechanical cell engineering, immunogenic cell death, whole-tumor-cell vaccination, iron oxide nanoparticles, cancer immunoengineering, bladder cancer

## Abstract

Whole-cell cancer vaccines can provide broad tumor-antigen repertoires, but their efficacy depends on manufacturing methods that inactivate tumor cells while preserving immunogenic danger signals. Here we show that laser-induced vapor nanobubbles (VNB) generated from dextran-coated iron oxide nanoparticles provide a drug-free photomechanical route to engineer immunogenic whole-cell vaccines against bladder cancer. Using MB49 murine urothelial carcinoma cells, we established VNB conditions that induced rapid tumor-cell killing together with calreticulin exposure and extracellular ATP and HMGB1 release. Rapid cell killing and calreticulin exposure were also reproduced in the molecularly distinct UPPL1541 bladder cancer model, supporting the generalizability of the early photomechanical response across bladder cancer cell lines. Vaccination with VNB-killed MB49 cells reduced tumor incidence and growth after subcutaneous challenge and enhanced splenocyte-mediated antitumor activity against MB49 cells. In an orthotopic bladder cancer model, vaccination prevented MRI-detectable tumor formation and was associated with marked early bladder enrichment of CD8⁺ T cells and inflammatory monocytes. A single administration of the VNB-generated vaccine preparation showed no evident local or systemic toxicity during seven days of follow-up. These findings provide proof of concept for an externally triggered, drug-free manufacturing approach in which short laser irradiation induces rapid photomechanical killing of tumor cells on demand, generating immunogenic whole-tumor-cell vaccine preparations for further evaluation in therapeutic, post-resection and tumor-derived settings.

## INTRODUCTION

Bladder cancer (BC) remains a major global health burden, ranking among the ten most common malignancies worldwide. According to the latest GLOBOCAN estimates, approximately 635,000 new cases and 228,000 deaths occurred worldwide in 2024, ranking bladder cancer as the eighth most commonly diagnosed cancer and the 13^th^ leading cause of cancer-related death globally.^1^ Clinically, the disease is divided into muscle-invasive bladder cancer (MIBC), which carries a high risk of dissemination, and non-muscle-invasive bladder cancer (NMIBC), which represents most newly diagnosed cases. Although NMIBC is usually associated with better initial survival, it behaves as a recurrent field disease: 60-70% of patients relapse after treatment, increasing the likelihood of disease progression, treatment-related morbidity and long-term healthcare costs.^2^ Thus, the central challenge in NMIBC is not only removal of visible lesions, but prevention of tumor re-emergence after intervention. Current NMIBC management is built around transurethral resection of bladder tumor (TURBT), followed by risk-adapted intravesical treatment. Local chemotherapy with agents such as mitomycin C or gemcitabine can reduce recurrence, but it is constrained by local toxicity and does not generate durable tumor-specific immune memory.^2^ Bacille Calmette-Guerin (BCG), a live attenuated form of *Mycobacteriumbovis*, remains the standard intravesical therapy for intermediate- and high-risk NMIBC because it activates innate and adaptive immune mechanisms in the bladder.^3^ However, BCG can cause substantial local inflammation, urinary symptoms and, in rare cases, disseminated BCG infection; moreover, between 30-50% of patients fail to respond and relapse after treatment.^4^ These limitations have renewed interest in prophylactic immune strategies that reinforce antitumor immunity after tumor resection. In this direction, BCG but also other mycobacteria-derived formulations such as RUTIVAC or purified protein derivative (PPD) skin testing have shown that a prime-boost strategy could enhance BCG-specific T-cell responses.^5–7^ Immune-checkpoint inhibitors targeting PD-1 or PD-L1 have also broadened the BC immunotherapy landscape, but their activity is often restricted by poorly inflamed or immunologically cold tumor microenvironments.^8^ Together, these observations motivate prophylactic strategies that combine broad tumor-antigen availability with immunogenic danger signaling.

Physical tumor-disruption approaches are particularly attractive in this context because they can couple tumor-cell damage with the release or exposure of immune-stimulatory signals. Phototherapies are particularly relevant for anatomically accessible tumors, including bladder lesions, because light can be delivered locally and with spatial control.^9,10^ Laser-based therapy is also considered less invasive than conventional surgery in selected settings,^11^ while nanoparticle-mediated photothermal therapy (PTT) can concentrate light-to-heat conversion at tumor sites more efficiently than many molecular absorbers.^12^ Beyond direct ablation, PTT has been increasingly linked to immunogenic cell death (ICD), a regulated mode of cell death in which dying tumor cells emit signals known as damage-associated molecular patterns (DAMPs) that promote both innate and adaptive antitumor immune responses.^10^ This connection is not simply a matter of delivering more heat. For example, in neuroblastoma, Prussian blue nanoparticle-mediated PTT revealed a thermal window in which intermediate temperatures induced stronger ICD-associated signaling than higher thermal doses.^13^ Similarly, temperature-dependent studies with functionalized gold nanoparticles showed that the mode of cell death can shift with photothermal intensity.^14^ These findings indicate that excessive or poorly confined heating may destroy antigens, damage surrounding tissue and weaken the quality of immune priming, which remains a major limitation of conventional PTT.^15^

The immunological consequences of tumor-cell death depend not only on the extent of cell killing, but also on the nature and timing of the DAMPs that are exposed or released.^16^ These endogenous signals can recruit and activate antigen-presenting cells (APCs), particularly dendritic cells (DCs), thereby providing an inflammatory context that supports tumor-antigen uptake and subsequent T-cell priming.^17^ Importantly, the detection of DAMP exposure or release *in vitro* is not sufficient on its own to establish immunogenic cell death. Consensus guidelines therefore recommend demonstrating functional antitumor protection in immunocompetent syngeneic vaccination models.^18^ Among the most informative ICD-associated readouts are cell-surface exposure of calreticulin (CALR) and extracellular release of both adenosine triphosphate (ATP) and high-mobility group box 1 (HMGB1). Specifically, CALR functions as a pro-phagocytic ’eat-me’ signal, ATP drives myeloid-cell recruitment and activation, and HMGB1 supports inflammatory antigen presentation.^19,20^ Photothermal nanomedicine has repeatedly been shown to engage this biology: gold nanoshell-mediated tumor-cell death activated inflammasome-associated responses,^21^ and subsequent studies combined photothermal damage with systemic immune activation or immune effector strategies in melanoma and other tumor models.^22,23^ Polypyrrole-based photothermal nanoparticles further demonstrated that nanomaterial morphology can influence immunological responses,^24^ while PTT targeting breast cancer stem-like populations reduced metastatic progression.^25^ More recently, magnetic nanoparticle platforms have been engineered to synergize macrophage polarization with ICD-associated tumor damage.^26^ Together, these studies establish PTT as an immunologically active modality, but also highlight the need for physical regimes that decouple lethal tumor damage from uncontrolled bulk heating.

A central limitation of conventional PTT is that increasing thermal dose to achieve efficient tumor-cell killing can promote heat diffusion beyond the immediate target, causing nonspecific tissue damage and potentially compromising temperature-sensitive cellular components. This motivates the development of photothermal regimes in which absorbed optical energy is converted into highly localized mechanical effects rather than sustained bulk heating. Laser-induced vapor nanobubbles (VNBs) offer such a regime. Upon irradiation with short, typically nanosecond, laser pulses, photothermal structures undergo rapid heating that transiently vaporizes the surrounding aqueous medium and generates a VNB. Bubble expansion and collapse produce pressure waves, fluid motion and localized mechanical stresses capable of permeabilizing or disrupting nearby biological membranes while minimizing heat diffusion into the surrounding environment.^27,28^ By tuning the optical and nanomaterial parameters, VNB-mediated membrane disruption can range from transient and reversible permeabilization to irreversible mechanical damage. This tunability has been exploited in photoporation to overcome different cellular barriers, including the plasma membrane, endosomal membrane and nuclear envelope, enabling controlled intracellular delivery of otherwise membrane-impermeable macromolecules.^29^ While mostly applied in an *in vitro* setting, the application of VNBs is also being considered for *in vivo* applications. In accessible tissues, plasmonic nanobubbles have been exploited for intraoperative detection and elimination of residual microtumors,^30^ and laser-induced VNBs have safely ablated vitreous opacities *in vivo*.^31^ These applications illustrate the central advantage of VNBs: the disruptive effect can be tuned through optical and nanomaterial parameters, ranging from transient membrane permeabilization to irreversible mechanical disruption.

This tunable disruption is especially attractive for whole-tumor-cell vaccination, a strategy based on killed tumor cells that can present a broad repertoire of tumor antigens rather than a single predefined target.^32,33^ This concept is particularly relevant for NMIBC, where relapse can arise from residual tumor cells and antigenically diverse clones that remain after local intervention. However, antigen breadth alone is unlikely to be sufficient. The vaccine substrate must be efficiently inactivated while retaining whole-cell-derived antigenic material and, at the same time, provide danger signals capable of promoting immune recognition.^32,33^ VNB-mediated disruption is well suited to address these requirements because it enables rapid and externally controlled on-demand tumor-cell killing through localized photomechanical damage, without relying on a cytotoxic drug or sustained bulk heating. Moreover, because the treatment disrupts rather than selectively targets a defined antigenic component, the resulting vaccine material retains the complexity of the whole tumor-cell source while potentially coupling this antigenic breadth to the exposure and release of immunogenic danger signals. An important precedent for this concept was provided by Ramon et al., who showed that nanosecond pulsed-laser irradiation of endogenous melanin-containing melanosomes in B16-F10 melanoma cells generated VNBs capable of inducing spatially confined mechanical cell killing together with the exposure and release of ICD-associated DAMPs, including CALR, ATP and HMGB1.^34^ These findings suggested that VNB-mediated mechanical disruption itself could generate an immunogenic form of tumor-cell damage. However, reliance on endogenous melanosomes inherently restricts this approach to highly pigmented cells. We therefore asked whether this phenomenon could be generalized to other tumor types by replacing the endogenous photothermal absorber with an externally supplied nanosensitizer. Among exogenous photothermal materials, iron oxide nanoparticles (IONPs) are particularly attractive because of their established use in biomedical applications, photothermal properties and generally favorable biodegradability and biocompatibility profiles.^35–37^ We selected dextran-coated IONPs because the polymer coating additionally provides aqueous colloidal stability and reduces nonspecific particle-associated cytotoxicity,^38^ while maintaining an externally controllable photothermal sensitizer for VNB generation. This provides a modular strategy in which VNB-mediated photomechanical killing is no longer dependent on the presence of an endogenous optical absorber.

Here, we use VNBs generated from dextran-coated IONPs as a drug-free photomechanical strategy to engineer immunogenic whole-tumor-cell vaccines against bladder cancer (Fig. 1). The conceptual workflow envisages using tumor-cell material collected after bladder tumor resection as an antigenic source for vaccine preparation (Fig. 1a). To establish experimental proof of concept, we used MB49 murine urothelial carcinoma cells for vaccine generation and subsequent evaluation in syngeneic vaccination–challenge models (Fig. 1b). Following incubation with IONPs, tumor cells are exposed to 532 nm nanosecond pulsed-laser irradiation, triggering VNB expansion and collapse and localized photomechanical disruption. We first establish the nanoparticle dose, incubation time and irradiation conditions required for efficient tumor-cell killing, and then characterize ICD-associated signals, including increased CALR positivity in dying/dead cells and extracellular ATP and HMGB1 release. We subsequently evaluate the protective activity of the resulting vaccine in subcutaneous and orthotopic bladder tumor challenge models, together with systemic antitumor activity, early bladder immune infiltration and short-term tolerability. This approach links whole-tumor-cell antigenic material with externally triggered on-demand photomechanical killing and immunogenic danger signaling, providing a foundation for further evaluation in tumor-derived and post-resection vaccination settings.

**Fig. 1:**
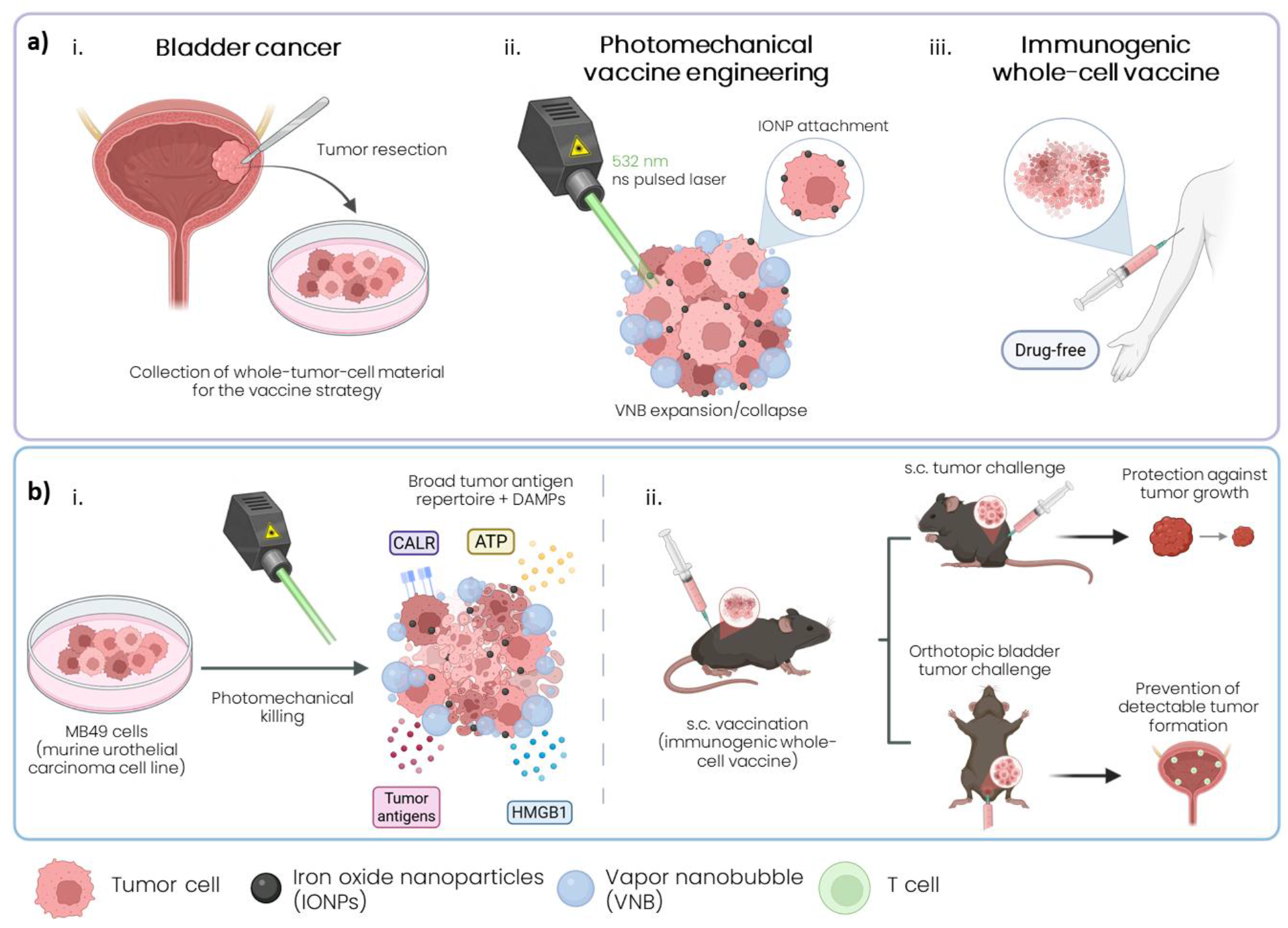
Photomechanical engineering of an immunogenic whole-cell vaccine for bladder cancer prophylaxis. a, Conceptual workflow for vaccine preparation from resected bladder tumor material. (i) Collection of tumor cells following bladder tumor resection as an antigenic source. (ii) Incubation with dextran-coated iron oxide nanoparticles (IONPs), followed by 532 nm nanosecond pulsed-laser irradiation. Rapid nanoparticle heating generates vapor nanobubbles (VNBs), whose expansion and collapse induce localized photomechanical disruption and tumor-cell killing. (iii) The resulting immunogenic whole-cell preparation is intended for administration as a drug-free cancer vaccine. b, Experimental implementation using MB49 murine urothelial carcinoma cells. (i) IONP-mediated VNB generation induces photomechanical killing, producing whole-tumor-cell vaccine material accompanied by ICD-associated damage-associated molecular patterns (DAMPs), including surface calreticulin (CALR) exposure and extracellular release of adenosine triphosphate (ATP) and high-mobility group box 1 (HMGB1). (ii) Subcutaneous vaccination with VNB-killed MB49 cells precedes challenge with viable MB49 cells in either a subcutaneous or an orthotopic bladder model. The outcomes illustrated are protection against subcutaneous tumor growth and prevention of detectable orthotopic bladder tumor formation, with T-cell infiltration represented in the bladder. Panel a depicts a prospective tumor-derived application; panel b summarizes the cell-line-based prophylactic strategy evaluated in this study. s.c., subcutaneous.

## RESULTS

### Optimization of IONP-mediated VNB generation and cell killing

To establish the experimental conditions required for VNB-mediated killing of bladder tumor cells, we first characterized dextran-coated IONPs as photothermal nanosensitizers for pulsed-laser activation. Dynamic light scattering analysis showed that the IONPs had a hydrodynamic diameter of 498 ± 21 nm (Fig. 2a-i), a ζ-potential of −11 ± 1 mV (Fig. 2a-ii), and a polydispersity index of 0.25 ± 0.03 (Fig. 2a-iii), consistent with a moderately dispersed colloidal formulation suitable for laser-induced VNB generation. We next determined the laser-fluence threshold required to generate VNBs around the IONPs using a custom pulsed-laser irradiation setup coupled to optical microscopy (Supplementary Fig. S1). Phase-contrast microscopy during nanosecond pulsed-laser irradiation at 532 nm showed the appearance of transient VNBs at 0.16 J/cm², with more consistent and readily detectable bubble formation from 0.39 J/cm² onwards (Fig. 2b). These results confirmed that dextran-coated IONPs can act as efficient nanosensitizers for VNB generation under short-pulse laser irradiation. Having established the minimal fluence for VNB-generation, we then optimized the treatment conditions in MB49 murine urothelial carcinoma cells. Cells were incubated with IONPs (0.1 mg/mL) for 10, 20 or 30 min, washed to remove non-associated particles, and exposed to pulsed-laser irradiation at increasing fluences (Fig. 2c). CellTiter-Glo measurements revealed a fluence-dependent decrease in MB49 viability, with limited cytotoxicity at the lowest fluences and a marked loss of viability at higher fluences (Fig. 2d). The effect was also influenced by nanoparticle incubation time, with 20 min of IONP incubation followed by irradiation at fluences ≥ 0.39 J/cm² providing robust tumor-cell killing while maintaining a practical treatment window for subsequent ICD experiments.

**Fig. 2:**
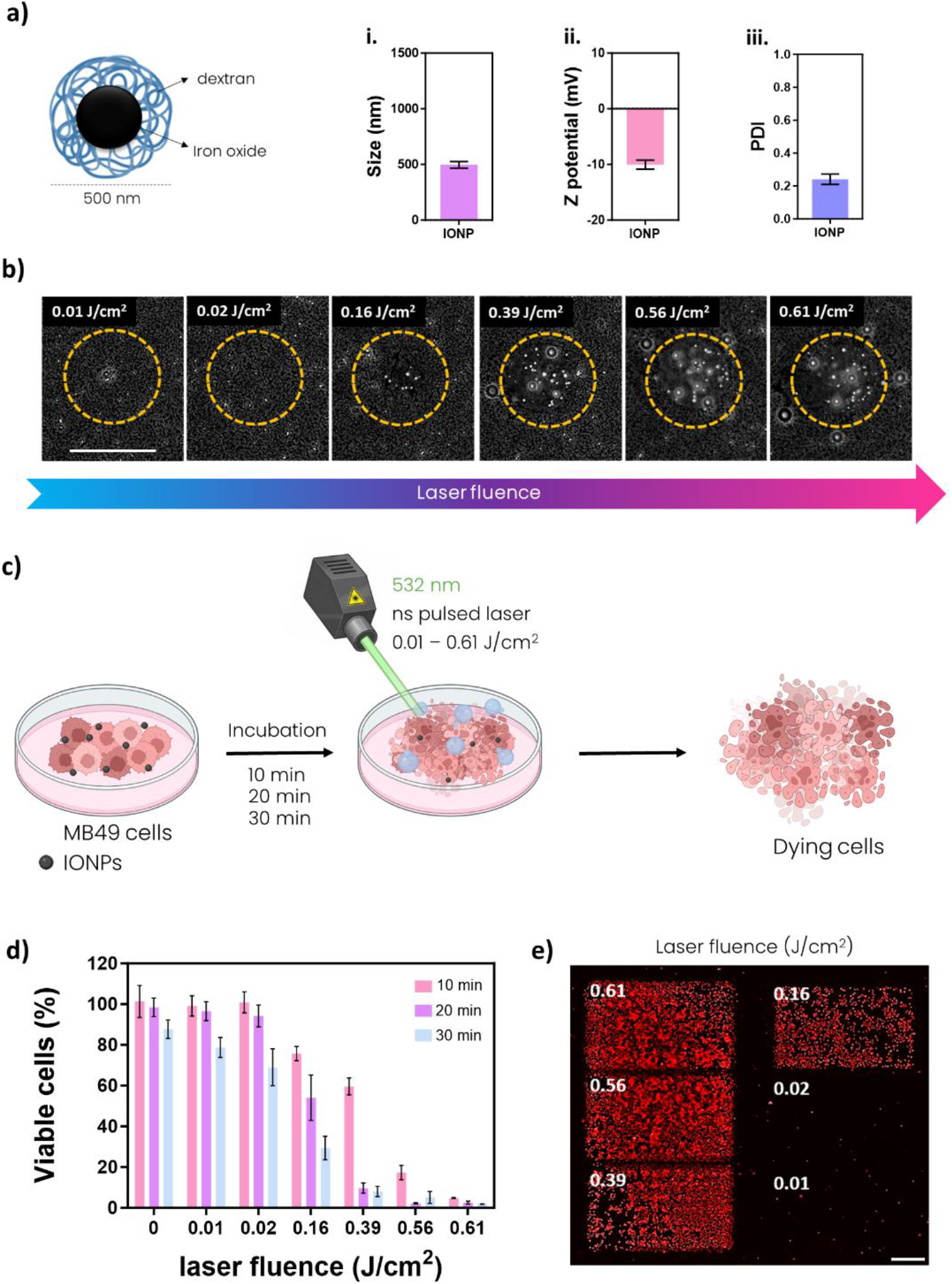
Optimization of VNB-mediated MB49 tumor-cell killing. a, Schematic and physicochemical characterization of dextran-coated iron oxide nanoparticles (IONPs). Hydrodynamic diameter, ζ-potential and polydispersity index were measured by dynamic light scattering. b, Representative phase-contrast images showing vapor nanobubble formation after pulsed-laser irradiation of IONPs at increasing laser fluences. The dashed yellow circles indicate the irradiated regions. The scale bars represent 100 µm. c, Schematic of the *in vitro* irradiation workflow: MB49 cells were incubated with IONPs and exposed to pulsed laser irradiation to induce VNB-mediated cell death. d, MB49 cell viability after IONP incubation (0.1 mg/mL) for 10, 20 or 30 min followed by irradiation at the indicated laser fluences, quantified using CellTiter-Glo. Data are shown as mean ± s.d.; n = 3. e, Representative fluorescence image of EthD-positive dead cells after irradiation at the indicated laser fluences. The scale bars represent 500 µm.

Consistent with the viability measurements, fluorescence imaging of ethidium homodimer-positive cells showed extensive membrane-compromised cell death after irradiation under the optimized VNB condition (Fig. 2e). Based on these results, 20 min IONP incubation and irradiation at 0.39 J/cm² were selected for all subsequent assays evaluating VNB-induced ICD.

Subsequently, we investigated whether VNB-mediated MB49 cell killing exhibited hallmarks of ICD. To this end, we first analyzed the kinetics and modalities of cell death induced by the different treatments. The cell-death populations evaluated by flow cytometry are schematically represented in Fig. 3a and were defined according to Annexin V–FITC and propidium iodide (PI) staining as viable, early apoptotic, late apoptotic or necrotic cells, following the gating strategy shown in Supplementary Fig. S2. Four experimental groups were compared throughout these assays: non-treated MB49 cells (NTC), cells incubated with IONPs but not irradiated (IONPs), cells incubated with IONPs and exposed to pulsed-laser irradiation to induce VNBs (VNBs), and doxorubicin-treated cells (DOX), which were included as a pharmacological positive control for ICD in subsequent experiments. For this, we first optimized the DOX killing condition in MB49 cells. Cells were incubated with increasing DOX concentrations for 24 h, and 120 µM DOX was selected because it reproducibly induced robust cell death 24 h after treatment (Supplementary Fig. S3a). Cell viability was quantified at 2, 4 and 6 h after treatment using two complementary approaches: CellTiter-Glo metabolic activity measurements and Annexin V–FITC/PI flow cytometry (Fig. 3b-i and 3b-ii, respectively). Both assays showed a rapid and marked decrease in MB49 cell viability after VNB treatment, whereas non-treated cells and IONP-treated cells remained largely viable and DOX induced only a modest reduction in viability within this early time window (Fig. 3b). Consistently, flow-cytometry analysis of the cells recovered after treatment showed a pronounced increase in the late-apoptotic population, while early-apoptotic and necrotic populations remained comparatively low across the analyzed time points (Fig. 3c). Because cells that underwent immediate VNB-induced lysis or extensive fragmentation would not be recovered for flow-cytometry analysis, these measurements describe the cell-death phenotype of the remaining analyzable cell population rather than the complete spectrum of VNB-induced killing. Notably, immediate post-treatment EthD-2 imaging showed extensive membrane-compromised cell death with numerous dead cells remaining attached after irradiation (Fig. 2e, and Fig. S3b-ii). Thus, VNB treatment rapidly compromises MB49 cell viability, with late-apoptotic cells becoming prominent among the cellular fraction recovered for subsequent flow-cytometry analysis.

**Fig. 3:**
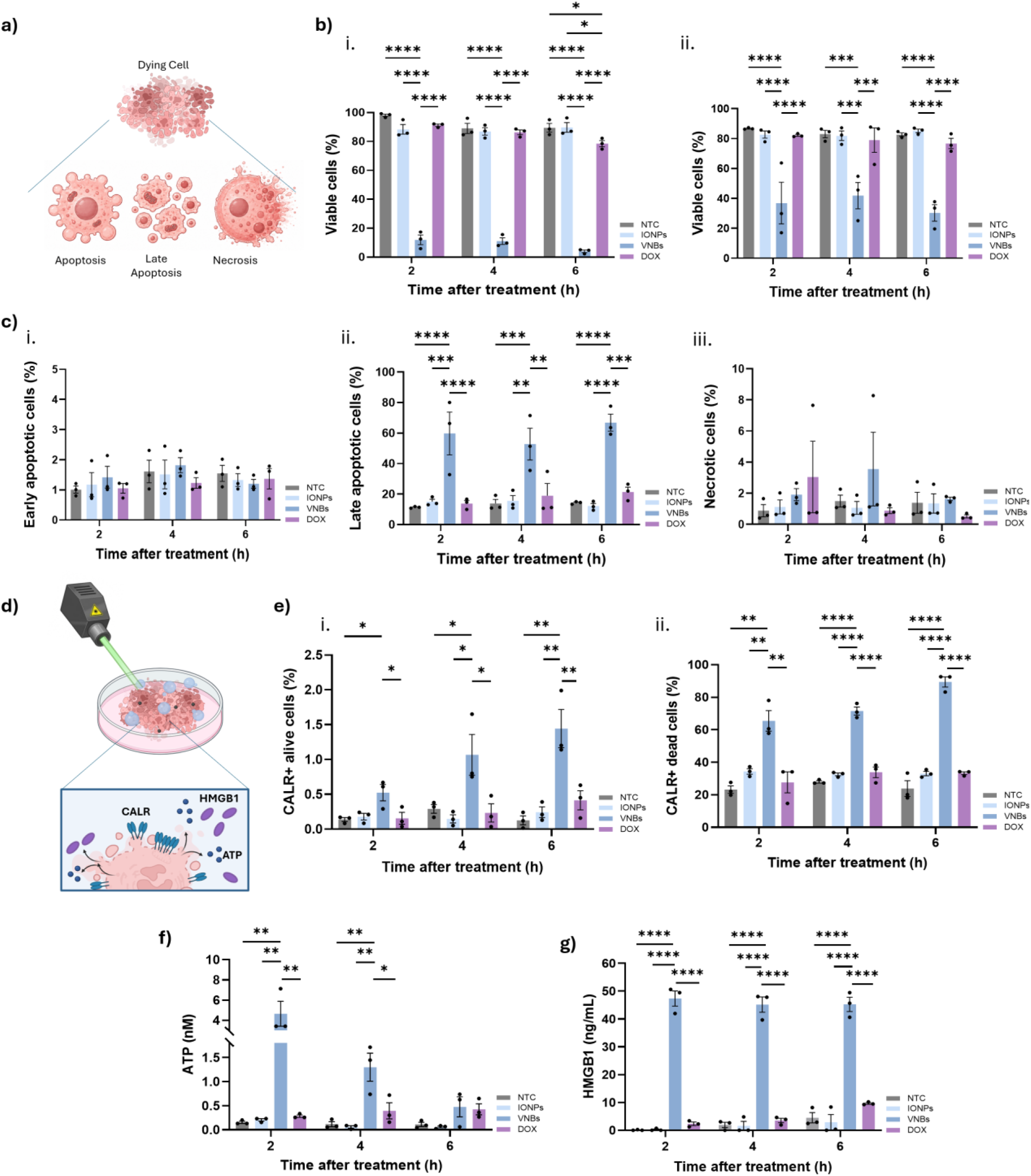
VNB-mediated mechanical disruption induces hallmarks of ICD in MB49 cells. a, Schematic representation of the cell-death modalities evaluated by Annexin V–FITC and propidium iodide staining: viable, early apoptotic, late apoptotic and necrotic cells. b, Time-course analysis of MB49 cell viability at 2, 4 and 6 h after treatment, determined by (i) CellTiter-Glo metabolic viability assay and (ii) Annexin V–FITC/PI flow cytometry, following the gating strategy shown in Supplementary Fig. S2. c, Time-course quantification of MB49 cell-death modalities by Annexin V–FITC/PI flow cytometry for (i) early apoptotic, (ii) late apoptotic and (iii) necrotic cells. d, Schematic representation of VNB-induced tumor-cell disruption and release or exposure of immunogenic damage-associated molecular patterns (DAMPs), including ATP, HMGB1 and calreticulin (CALR). e, Time-course quantification of (i) CALR-positive viable cells and (ii) CALR-positive dying/dead cells at 2, 4 and 6 h after treatment. f, Extracellular ATP release measured in culture supernatants at 2, 4 and 6 h after treatment. g, Extracellular HMGB1 release measured in culture supernatants at 2, 4 and 6 h after treatment. Experimental groups include non-treated cells (NTC), IONP-treated cells (IONPs), VNB-treated cells (VNBs) and doxorubicin-treated cells (DOX). Doxorubicin was used as a positive control for immunogenic cell death. Data are shown as mean ± SEM.; individual points indicate independent biological replicates; n = 3. Statistical significance was determined by two-way ANOVA followed by Tukey’s multiple-comparisons test. Statistically significant differences are shown as: \**p* < 0.05, \*\**p* < 0.01, *** *p* < 0.001, \*\*\*\**p* < 0.0001.

Next, we quantified the emission and exposure of key ICD-associated DAMPs, focusing on CALR, ATP and HMGB1, which respectively provide pro-phagocytic, chemoattractant and inflammatory signals to APCs (Fig. 3d). CALR cell-surface exposure was assessed at 2, 4 and 6 h after treatment by spectral flow cytometry and quantified separately within viable and dying/dead-cell populations using Annexin V–FITC and PI staining. VNB-treated cells showed a clear increase in CALR-positive viable cells over time compared with untreated and IONP-treated controls (Fig. 3e-i). In parallel, VNB treatment also increased the fraction of CALR-positive dying/dead cells (Fig. 3e-ii), again exceeding the response observed in DOX-treated cells across the evaluated time points, and indicating that VNB-mediated mechanical disruption was particularly efficient at promoting this early “eat-me” signal. Because CALR staining was performed without permeabilization, CALR positivity in viable cells primarily reflects surface-exposed CALR. In contrast, interpretation within the dying/dead population requires caution, as loss of plasma-membrane integrity in PI-positive cells may allow antibody access to intracellular CALR and contribute to the baseline CALR signal observed in untreated, IONP-treated and DOX-treated controls. Despite this limitation, VNB treatment produced a marked increase in CALR-positive dying/dead cells compared with the corresponding control groups. The gating strategy used to define viable, dying/dead and CALR-positive populations is provided in Supplementary Fig. S4a.

Extracellular ATP, a soluble DAMP involved in myeloid-cell recruitment and immune activation, was rapidly released after VNB treatment. ATP concentrations were calculated from a standard calibration curve generated in parallel (Supplementary Fig. S4b). ATP levels in the culture supernatant increased at early time points after irradiation and remained elevated across the analyzed time-course relative to the negative controls (Fig. 3f). Notably, VNB-treated cells released higher ATP levels than DOX-treated cells at 2, 4 and 6 h time points, showing that VNB-mediated killing elicited an earlier release of ATP than the pharmacological ICD positive control.

Finally, we evaluated the release of HMGB1, a late ICD-associated DAMP linked to inflammatory activation and antigen presentation. HMGB1 concentrations were derived from the corresponding assay calibration curve (Supplementary Fig. S4c). VNB-treated MB49 cells released higher levels of extracellular HMGB1 than the negative control cells across the analyzed time points (Fig. 3g). Overall, VNB treatment triggers the robust exposure and release of several hallmarks associated with ICD, including CALR exposure and ATP, and HMGB1 release. Functional vaccination–challenge experiments were subsequently used to determine whether these signals were accompanied by protective antitumor immunity.

To further evaluate the persistence and evolution of ICD-associated DAMPs, we extended the analysis to 24 h after treatment (Supplementary Fig. S5). At this later time point, DOX, used as the pharmacological positive control for ICD, produced the highest CALR positivity among viable cells and the highest extracellular ATP and HMGB1 levels (Supplementary Fig. S5). Importantly, despite being triggered by a brief irradiation event, VNB treatment still resulted in significantly increased ICD-associated signals 24 h after photomechanical perturbation compared with untreated controls. In particular, VNB-treated samples maintained a pronounced increase in CALR-positive dying/dead cells relative to untreated and IONP-treated controls (Supplementary Fig. S5a-ii), together with elevated extracellular ATP and HMGB1 levels. Overall, these results indicate that the rapid DAMP response initiated by VNB-mediated disruption is followed by persistent ICD-associated signals that remain detectable 24 h after irradiation.

To determine whether the cellular response to VNB treatment was restricted to the MB49 model, we repeated the early time-course analysis in UPPL1541 cells, a murine bladder cancer cell line that retains molecular characteristics of the luminal-like subtype of human urothelial carcinoma. VNB treatment induced a rapid and sustained reduction in metabolic viability, with only approximately 20% of cells remaining viable from 2 h onwards, whereas untreated, IONP-treated and DOX-treated cells retained substantially higher metabolic activity throughout the evaluated period (Supplementary Fig. S6i). Consistent with this loss of viability, VNB treatment caused a progressive accumulation of late-apoptotic cells, increasing from approximately 45% at 2 h to more than 70% at 6 h after treatment (Supplementary Fig. S6ii). In contrast, DOX induced only a modest increase in late apoptosis during this early time window. VNB treatment also promoted a marked time-dependent increase in CALR-positive dying/dead UPPL1541 cells. CALR positivity increased from approximately 20% at 2 h to more than 60% at 4 h and approximately 75% at 6 h, whereas untreated and IONP-treated cells showed minimal CALR exposure (Supplementary Fig. S6iii). DOX produced a smaller and more delayed increase in CALR-positive dying/dead cells. Thus, VNB-mediated photomechanical disruption reproduced rapid cell killing, late-apoptotic progression and exposure of the pro-phagocytic CALR signal in a molecularly distinct bladder cancer cell line, supporting the robustness of the response beyond the MB49 model.

### Vaccination with VNBs-treated MB49 cells protects against subcutaneous tumor challenge

We next investigated whether the enhanced ICD profile induced by VNBs could translate into protective antitumor immunity *in vivo*. To this end, female mice were allocated to three groups: animals vaccinated subcutaneously with VNB-treated MB49 cells (VNB-based vaccine), animals vaccinated with DOX-treated MB49 cells (DOX-based vaccine), and unvaccinated controls (Unvaccinated). One week later, all animals were challenged with viable MB49 cells in the contralateral flank (Fig. 4a). This prophylactic vaccination–challenge setting allowed us to test whether killed tumor cells generated by VNBs could function as a syngeneic whole-cell vaccine. The optimized conditions used to generate DOX- and VNB-killed MB49 preparations with a dead-cell fraction above 90% are shown in Supplementary Fig. S3, including the DOX concentration-response analysis after 24 h of treatment (Supplementary Fig. S3a) and the high-throughput VNB-mediated cell-killing workflow after IONP loading and pulsed-laser irradiation (Supplementary Fig. S3b). Representative images of each group, with arrows and magnified insets indicating tumor formation, are shown in Fig. 4b. Tumors were apparent in control animals, whereas almost all VNB-based vaccinated mice remained tumor-free after challenge. The magnitude of protection was comparable to that observed in DOX-based vaccinated mice, supporting the capacity of VNBs-treated cells to induce functional antitumor immunity *in vivo*. Tumor development was markedly reduced in both vaccinated groups relative to unvaccinated controls. Quantification of tumor size showed that VNB-based vaccinated mice had significantly smaller tumors than control mice at the analyzed time points (Fig. 4c). Tumor incidence was 25% in the VNB-based vaccinated group (3/12 animals), 27.3% in the DOX-based vaccinated group (3/11 animals), and 100% in the unvaccinated group (12/12 animals) (Fig. 4d). To determine whether vaccination also enhanced splenocyte-mediated antitumor activity, splenocytes isolated from vaccinated or control mice were co-cultured with viable MB49 cells. Splenocytes from VNB-based vaccinated mice significantly reduced MB49-associated viability/proliferation after co-culture compared with splenocytes from unvaccinated control animals (Fig. 4e), supporting enhanced splenocyte-mediated antitumor activity following vaccination. A similar effect was observed in the DOX-based vaccinated group, consistent with the expected immunogenicity of this positive-control treatment.

**Fig. 4:**
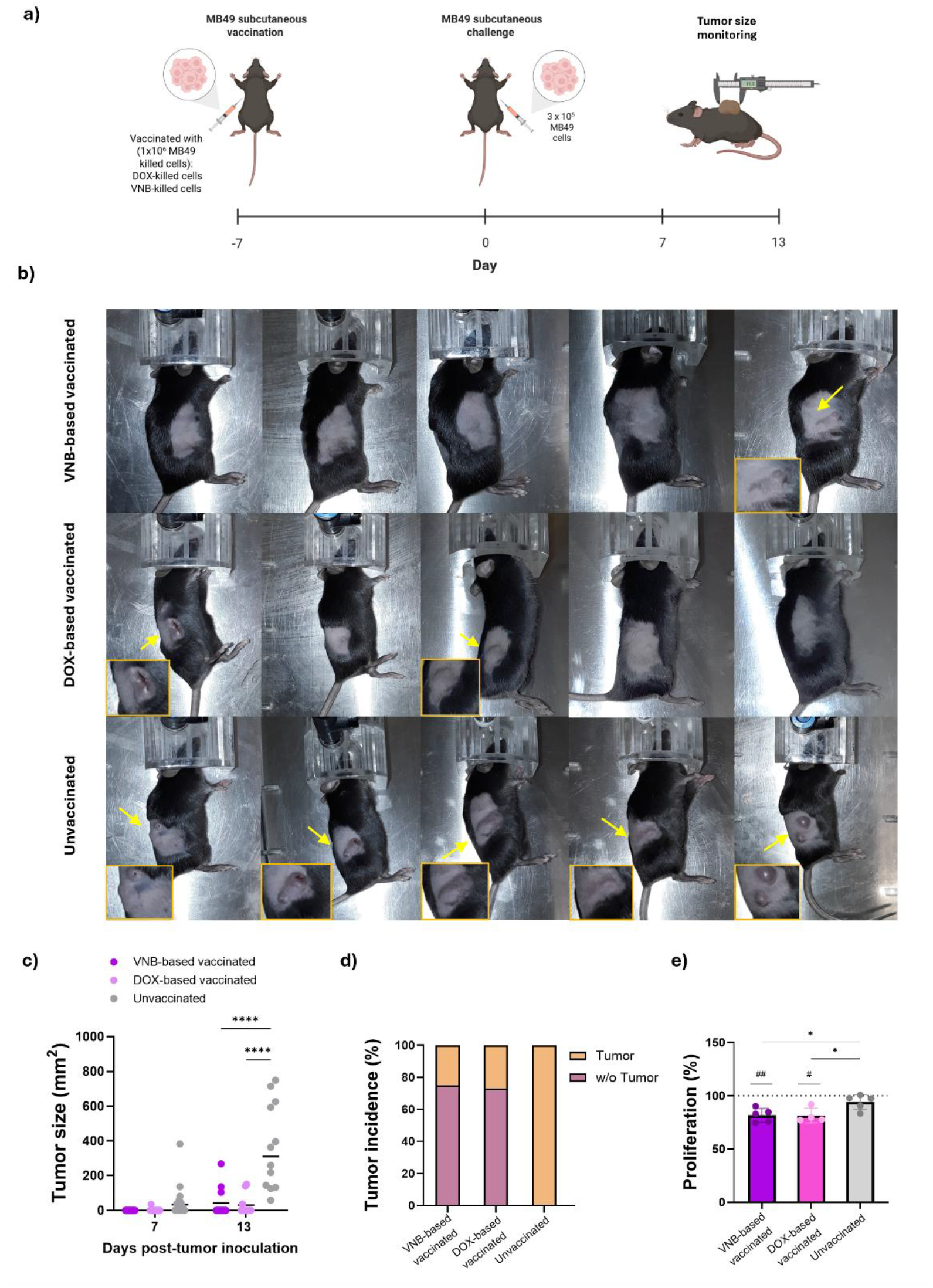
VNB-based vaccination protects mice against subcutaneous tumor challenge. a, Experimental design of the prophylactic subcutaneous vaccination–challenge model. Mice were vaccinated subcutaneously with 1 × 10⁶ dead MB49 cells generated by VNB or DOX treatment and challenged 7 days later with 3 × 10⁵ viable MB49 cells in the contralateral flank. Tumor growth was monitored by caliper measurement. b, Representative images of mice after subcutaneous tumor challenge. Arrows and magnified insets indicate visible tumor lesions. c-d, Tumor size (c) and tumor incidence (d) at the indicated time points after MB49 challenge in VNB-based vaccinated (n = 12) and DOX-based vaccinated (n = 11) mice, or unvaccinated mice (n = 12). Individual points represent individual mice, and horizontal lines indicate the mean value. One animal from the DOX-based vaccinated group was lost during the study because of an unrelated death. e, *In vitro* splenocyte-mediated antitumor activity assessed by co-culture with MB49 cells followed by MTT quantification of residual adherent MB49 cell proliferation. Proliferation was expressed relative to MB49 cells cultured alone, considered as 100% (dotted line). Data are shown as mean ± s.d.; individual points indicate splenocyte cultures from individual mice (VNB-based vaccinated, n = 5; DOX-based vaccinated, n = 4; Unvaccinated, n = 5). Tumor size in c) was analyzed by two-way repeated-measures ANOVA followed by Tukey’s multiple-comparisons test. \*\*\*\**p* < 0.0001. In e), differences relative to 100% proliferation were analyzed using one-sample t-tests followed by Holm–Šídák correction (^#^), whereas differences among experimental groups were analyzed by one-way ANOVA followed by Tukey’s multiple-comparisons test (*). \**p* < 0.05, \*\**p* < 0.01; #*p* < 0.05, ##*p* < 0.01.

Since sex can influence antitumor immune responses, we also evaluated the vaccination–challenge strategy in male mice (Supplementary Fig. S7). Representative images and tumor-size quantification confirmed that the protective effect of VNB-killed cells was preserved in males, as VNB- or DOX-based vaccinated mice showed reduced subcutaneous tumor growth compared with control animals. Having established that the vaccination effect was not restricted to female mice in the subcutaneous setting, subsequent orthotopic bladder cancer experiments were performed in females, in which transurethral catheterization and intravesical instillation can be performed more reproducibly owing to anatomical accessibility.

### VNB-engineered whole-cell vaccination shows a favorable safety profile

We next evaluated the short-term safety of the whole-cell vaccine preparations under the same conditions used in the vaccination experiments. Mice received a single subcutaneous injection of VNB-based vaccine, DOX-based vaccine, or DMEM alone as a vehicle control. General condition, body weight, local tolerability, organ morphology, and plasma biochemical parameters were evaluated for up to 7 days after administration (Fig. 5).

**Fig. 5:**
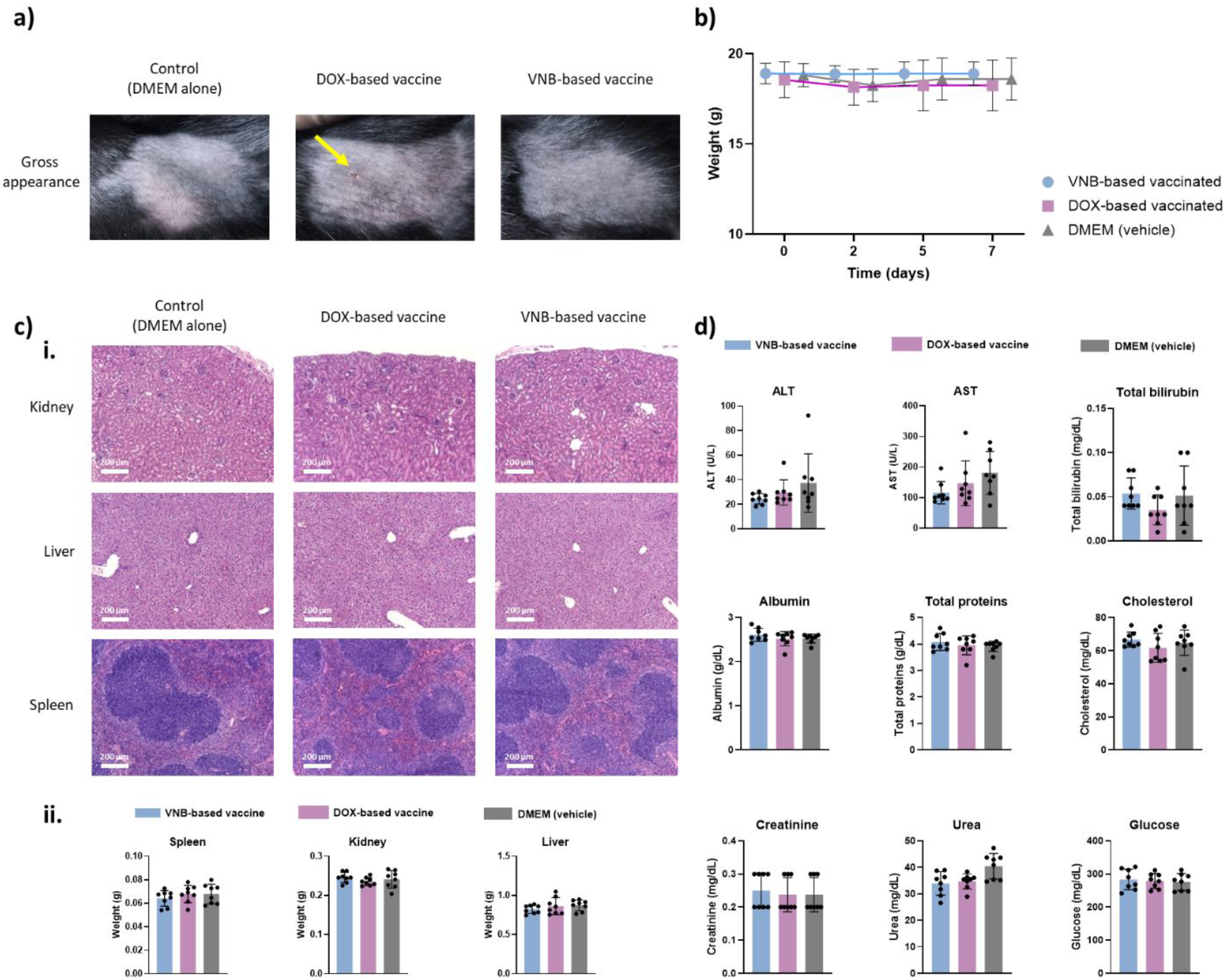
Short-term safety evaluation of VNB-engineered whole-cell vaccination. a, Representative gross images of the injection site 7 days after administration of vehicle (DMEM), DOX-based or VNB-based vaccines. Mild localized ulceration was observed in three of eight mice receiving DOX-treated cells (highlighted with a yellow arrow). b, Body weight measured before administration and at days 2, 5 and 7 after vaccination. c, Evaluation of major organs at day 7. (i) Representative H&E-stained sections of kidney, liver and spleen. Magnification, 10x objective. Scale bars, 200 μm. (ii) Absolute weights of the spleen, kidney and liver. d, Plasma biochemical analysis at day 7, including alanine aminotransferase (ALT), aspartate aminotransferase (AST), total bilirubin, albumin, total protein, cholesterol, creatinine, urea and glucose. Data are shown as mean ± s.d.; individual points represent individual mice; n = 8 mice per group. No significant differences were detected among groups.

Gross examination of the injection site at day 7 showed no evident skin alterations in mice receiving the VNB-based vaccine or DMEM alone, whereas mild localized ulceration or skin alteration was observed in three of eight mice receiving the DOX-based vaccine (Fig. 5a). Body weight remained stable throughout the observation period, with comparable values among the three experimental groups at days 0, 2, 5, and 7 (Fig. 5b). No treatment-associated weight loss or deterioration in general condition was observed, indicating the absence of overt systemic toxicity during the first week after administration.

To assess potential effects on major organs, the kidney, liver and spleen were collected at day 7 for histopathological evaluation and organ-weight determination. No treatment-associated histopathological alterations were identified in these organs, and absolute organ weights were comparable among the vaccination with VNB-killed and DOX-killed cells, and vehicle-control groups (Fig. 5c).

Plasma biochemical analysis further supported the absence of treatment-associated systemic toxicity (Fig. 5d). The hepatic injury markers alanine aminotransferase (ALT) and aspartate aminotransferase (AST), together with total bilirubin, were comparable among groups and showed no coordinated increase after administration of either vaccine preparation. Albumin and total protein concentrations were also maintained, indicating preservation of hepatic synthetic function. Similarly, plasma creatinine and urea concentrations showed no treatment-related changes indicative of impaired renal function. Cholesterol and glucose levels remained comparable among groups.

Together, these findings indicate that a single administration of either whole-cell vaccine preparation was well tolerated over the evaluated 7-day period, with no detectable effects on body weight, major-organ architecture, organ weights or plasma biochemical parameters. Gross examination at day 7 showed no evident injection-site alterations in mice receiving the VNB-based vaccine or vehicle control, whereas localized skin alterations were observed in three of eight mice receiving DOX-killed cells. Overall, the results support a favorable short-term systemic safety profile for both vaccine preparations, with no gross evidence of local alteration following administration of the VNB-based vaccine under the conditions evaluated.

### VNB-based vaccination prevents orthotopic bladder tumor formation

We next evaluated whether the protective effect observed in the subcutaneous challenge model could be reproduced in a clinically more relevant orthotopic bladder cancer setting. For this, female mice were vaccinated subcutaneously with VNBs- or DOX-based vaccines, or left unvaccinated, and challenged one week later by intravesical instillation of viable MB49 cells (Fig. 6a). Tumor burden was then assessed by MRI 21–24 days after intravesical tumor challenge and tumor volume was subsequently quantified from the MRI images. Notably, VNB-based vaccination prevented MRI-detectable orthotopic tumor formation in all vaccinated mice. Representative MRI images showed clear intravesical tumor masses in unvaccinated mice, whereas no detectable tumors were observed in the VNB-based vaccinated group (Fig. 6b-c). In the DOX-based vaccinated group, only one mouse developed a detectable tumor, with a tumor volume higher than those observed in unvaccinated tumor-bearing mice (Fig. 6b-c). Tumor incidence analysis quantified at the end of the experiment (euthanasia at day 30 after challenge) confirmed this effect, with no tumors detected in the VNB-based vaccinated group (0% incidence), while tumors were observed in all unvaccinated mice (100% incidence; Fig. 6d). Of note, one unvaccinated mouse did not show measurable tumor mass by MRI on day 21 after challenge, but bladder tumor was identified at necropsy, indicating 100% incidence in the unvaccinated group. Finally, DOX-based vaccination also reduced tumor incidence, with tumors detected in only 20% of mice (Fig. 6d). This finding extends the protection observed in the subcutaneous challenge model and supports the potential of VNB-induced ICD as a prophylactic whole-cell vaccination strategy for bladder cancer.

**Fig. 6:**
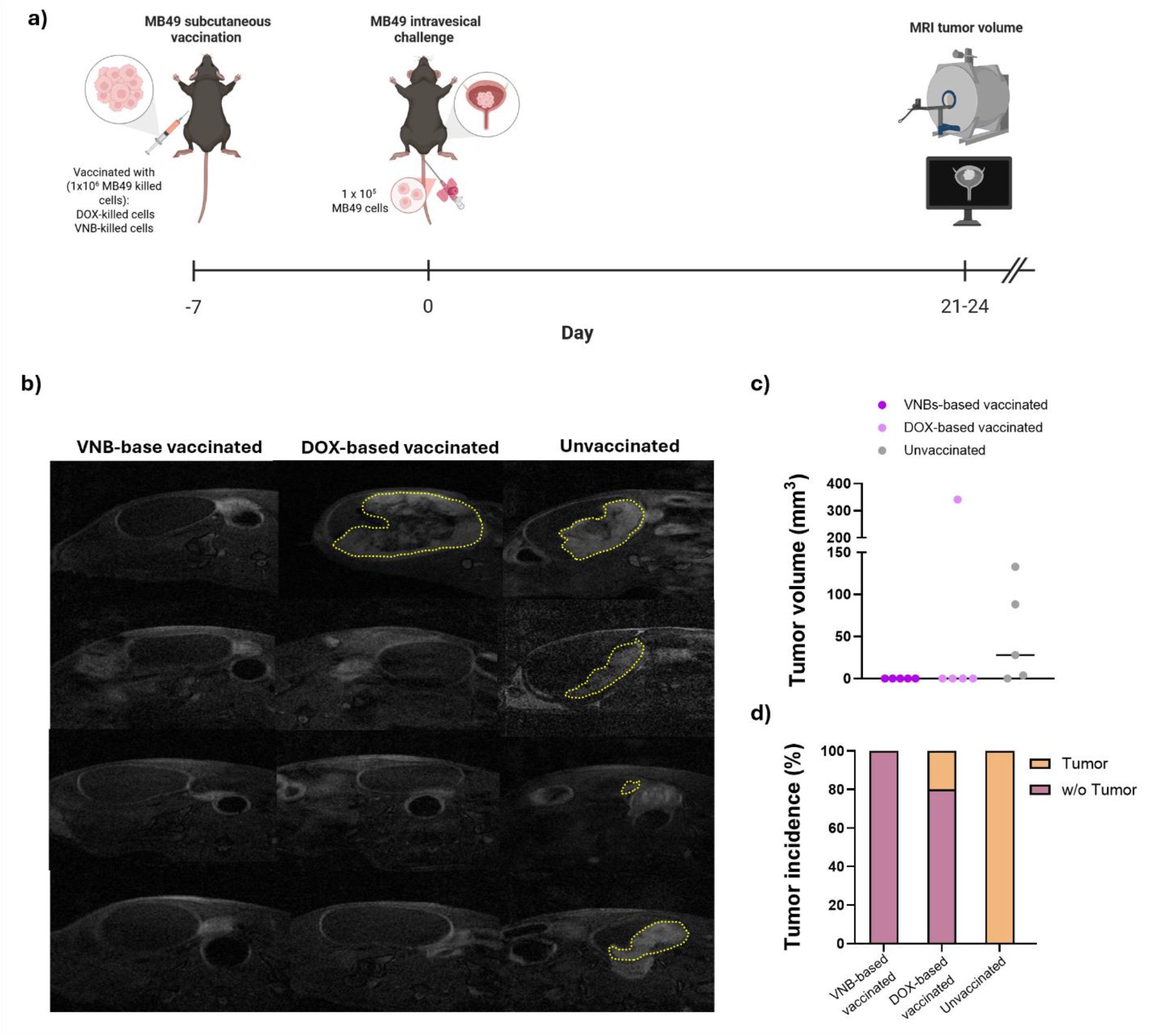
VNB-based vaccination prevents orthotopic bladder tumor formation. a, Experimental design of the prophylactic orthotopic bladder cancer model. Mice were vaccinated subcutaneously with 1 × 10⁶ VNB- or DOX-killed MB49 cells and challenged 7 days later by intravesical instillation of 1 × 10⁵ viable MB49 cells. Tumor burden was assessed by MRI 21-24 days after challenge. b, Representative axial MRI images of bladders from VNB-based vaccinated, DOX-based vaccinated and unvaccinated mice. Dashed outlines indicate detectable tumor areas. c, MRI-derived tumor volume in each experimental group. Individual points represent individual mice (n = 5) and horizontal lines indicate the mean value. Tumor volume was analyzed using the Kruskal–Wallis test followed by Dunn’s multiple-comparisons test. d, Tumor incidence in each experimental group including inspection in endpoint necropsy, expressed as the percentage of mice with or without bladder tumors.

### VNB-based vaccination enhances early immune infiltration in the bladder after orthotopic challenge

To investigate the immune changes associated with protection in the orthotopic model, we analyzed the early bladder immune landscape. For this, mice were vaccinated with VNB- or DOX-killed MB49 cells one week before challenge, and bladders were collected for flow-cytometry analysis two days after intravesical tumor challenge (Fig. 7a). Myeloid populations were identified using the gating strategy shown in Supplementary Fig. S8, whereas lymphoid populations were analyzed using the gating strategy shown in Supplementary Fig. S9. The antibody panels used for these analyses are listed in Supplementary Table S1.

**Fig. 7:**
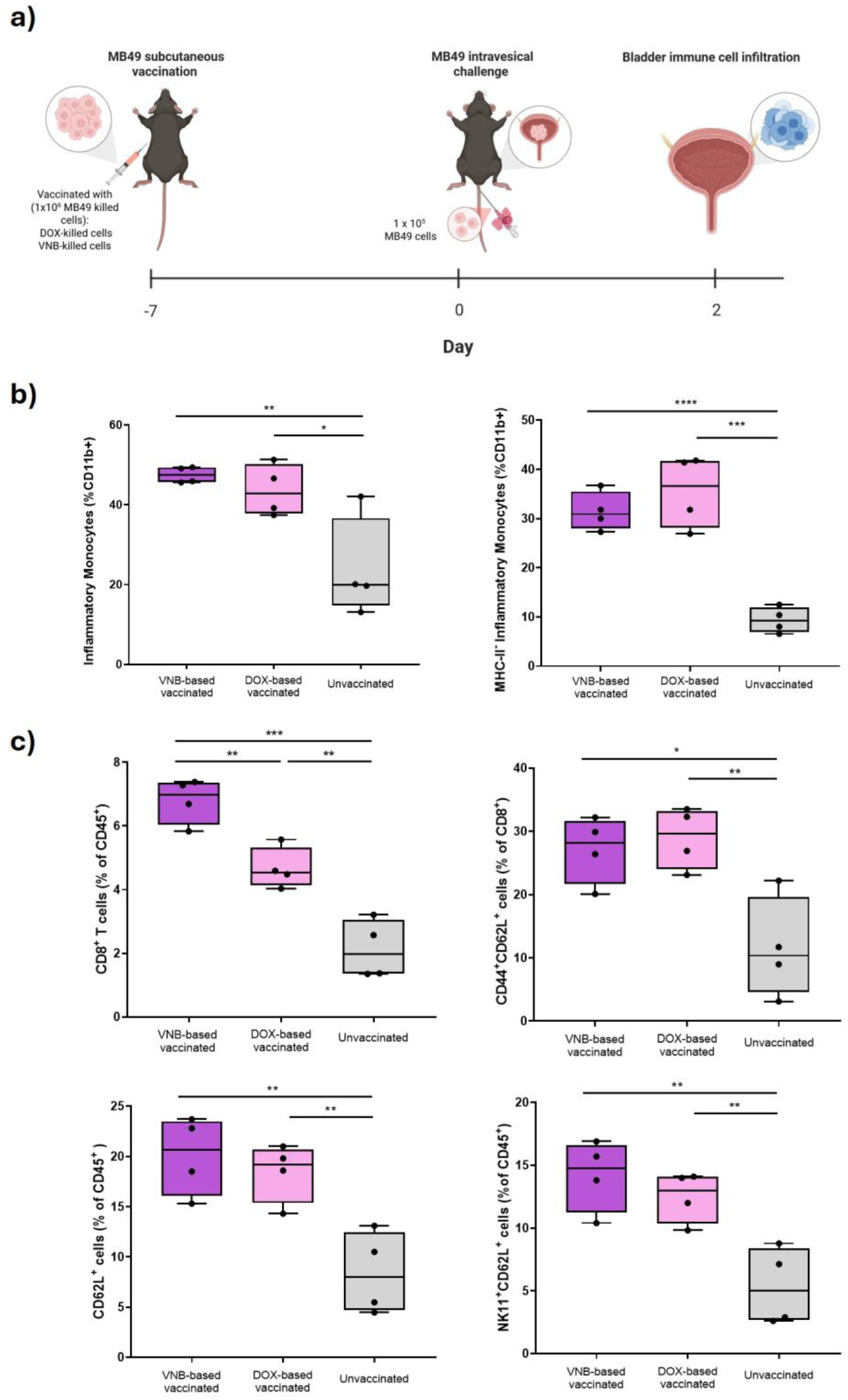
Vaccination enhances early immune infiltration and reshapes the bladder immune landscape after orthotopic challenge. a, Experimental design for early immune profiling. Mice were vaccinated subcutaneously with VNB- or DOX-killed MB49 cells, challenged 7 days later by intravesical instillation of viable MB49 cells, and bladders were collected 2 days after challenge for flow-cytometry analysis. b, Quantification of myeloid populations in the bladder, including inflammatory monocytes (CD11b⁺CD11c⁻Ly6G⁻Ly6C⁺) and MHC-II⁻ inflammatory monocytes (CD11b⁺CD11c⁻Ly6G⁻Ly6C⁺MHC-II⁻). c, Quantification of lymphoid immune populations infiltrating the bladder, including CD8⁺ T cells among CD45⁺ leukocytes, CD44⁺CD62L⁺ CD8⁺ T cells with a central memory-like phenotype, CD62L⁺ leukocytes and NK1.1⁺CD62L⁺ cells. Box plots show median, interquartile range and range; n = 4. Statistical significance was determined by two-sided Mann–Whitney U tests; *p < 0.05, **p < 0.01, ***p < 0.001.

We proceeded to analyses if vaccination induced a marked remodeling of the early bladder immune infiltrate. For this, we first focused on the myeloid compartment, as Ly6C⁺ monocytes represent an early innate immune population rapidly recruited to sites of tissue damage, infection or tumor implantation.^39,40^ Both VNB-based and DOX-based vaccinated mice showed increased infiltration of CD11b⁺CD11c⁻Ly6G⁻Ly6C⁺ monocytes, predominantly MHC-II⁻ inflammatory monocytes (IM), compared with unvaccinated controls, with no significant differences between the two vaccinated groups (Fig. 7b).

We next analyzed the lymphoid compartment. VNB-based vaccination induced a marked increase in CD8⁺ T-cell infiltration among CD45⁺ leukocytes compared with unvaccinated and DOX-based vaccinated mice (Fig. 7c). Vaccinated mice also showed increased frequencies of CD44⁺CD62L⁺ CD8⁺ T cells, a phenotype commonly associated with central memory-like T cells. In addition, both VNB- and DOX-based vaccines increased the percentages of CD62L⁺ leukocytes and NK1.1⁺CD62L⁺ cells in the bladder. Together, these findings indicate that whole-cell vaccination is associated with altered lymphoid composition in the bladder shortly after tumor challenge, with VNB-based vaccination producing a particularly pronounced enrichment of infiltrating CD8⁺ T cells. This immune remodeling occurred in the context of the functional antitumor protection observed in the vaccination–challenge experiments. The enrichment of CD8⁺ T cells is therefore compatible with a T-cell-associated component of the protective response, although the cytotoxic function and antigen specificity of these cells were not directly assessed.The CD62L profile provides additional information on the phenotype of the lymphoid response. CD62L, or L-selectin, is involved in lymphoid trafficking and homing, and CD44⁺CD62L⁺ expression is commonly associated with a central memory-like CD8⁺ T-cell phenotype. In this context, the increase in CD44⁺CD62L⁺ CD8⁺ T cells indicates enrichment of a memory-associated phenotypic population following vaccination. Although antigen specificity and functional recall capacity were not directly evaluated in this study, these data support a model in which vaccination promotes both early bladder infiltration by CD8⁺ T cells and enrichment of CD8⁺ T cells displaying a central memory-like phenotype.In addition to CD8⁺ T cells, VNB- and DOX-based vaccination increased the frequencies of CD62L⁺ leukocytes and NK1.1⁺CD62L⁺ cells in the bladder (Fig. 7c). These changes further indicate that whole-cell vaccination alters the composition and trafficking associated phenotype of the lymphoid compartment during the early response to intravesical tumor challenge.

## DISCUSSION

Our findings position VNB-mediated tumor-cell killing within the evolution of PTT from bulk heat-mediated ablation toward more controllable physical regimes of cell damage. Conventional nanoparticle-mediated PTT induces tumor-cell death through hyperthermia, membrane disruption, protein denaturation, DNA damage and, at high thermal doses, necrotic injury.^10^ As discussed above, PTT-induced immunogenicity depends not only on the extent of tumor-cell killing but also on the physical conditions under which cell damage is generated, with excessive heating potentially compromising the quality of the resulting immunogenic response.^13–15^ Photoporation and VNB-based approaches extend this concept by partially decoupling photothermal absorption from bulk heating. By tuning laser and nanomaterial parameters, membrane disruption can range from transient permeabilization for intracellular delivery to irreversible mechanical disruption and cell death.^41,42^ Mechanistically, VNBs arise from rapid local heating of photothermal structures after short laser pulses, producing bubble expansion and collapse that generate pressure waves, fluid jets and mechanical stresses while limiting heat diffusion.^29,41^ This mechanism was previously exploited using endogenous melanosomes in B16-F10 melanoma cells, where VNBs induced spatially controlled killing, DAMP exposure/release and membrane permeabilization.^34^ Here, we extend this principle to bladder cancer using exogenous dextran-coated IONPs, making the absorber externally defined, tunable and independent of tumor pigmentation. The use of dextran-coated IONPs is also translationally relevant, given the extensive biomedical history of IONPs in imaging, targeting, drug delivery and hyperthermia, including clinically approved or clinically investigated formulations.^35–37^

In this study, we first established the conditions required for controlled VNB formation and MB49 killing. The irradiation setup is shown in Supplementary Fig. S1, while Fig. 2 summarizes IONP characterization, fluence-dependent VNB formation and optimization of cell killing. Under the selected condition—20 min IONP incubation followed by irradiation at 0.39 J/cm²—VNB treatment induced robust MB49 cell death, whereas IONPs alone showed limited toxicity (Fig. 2; Supplementary Fig. S2). VNB-mediated killing was not merely cytotoxic: it caused a rapid loss of metabolic activity and flow-cytometry-defined viability (Fig. 3b), induced a predominantly late-apoptotic cell-death profile (Fig. 3c; Supplementary Fig. S2), and although early apoptosis drives initial DAMP trafficking,^43^ progression toward a prominent late-apoptotic state is expected to favor the extracellular release of nuclear and cytoplasmic danger signals. Importantly, irrespective of the relative contribution of different cell-death modalities, VNB treatment promoted the early exposure or release of CALR, ATP and HMGB1 (Fig. 3d–g; Supplementary Fig. S4), with the 24 h DAMP analysis summarized in Supplementary Fig. S5. These DAMPs are functionally relevant because they contribute to the inflammatory context associated with immunogenic cell death and can support subsequent antigen uptake and presentation by antigen-presenting cells.^16,18^

The reproducibility of the early VNB response in UPPL1541 cells further strengthens the biological relevance of the platform. UPPL1541 was derived from a genetically engineered luminal-like bladder tumor and retains epithelial and molecular features that more closely resemble human urothelial carcinoma and normal murine urothelium than MB49 cells, which exhibit a more mesenchymal, fibroblast-like transcriptional phenotype.^44^ Despite these differences, VNB treatment caused rapid loss of UPPL1541 viability, progressive late-apoptotic cell death and a pronounced increase in CALR exposure among dying/dead cells. This response was substantially faster and stronger than that induced by DOX during the same early time window. Previous work has shown that nanoparticle-mediated phototherapy can promote CALR redistribution towards the plasma membrane in UPPL cells.^45^ Our findings extend those observations by demonstrating quantitative and time-dependent CALR exposure following a drug-free, predominantly photomechanical treatment. Together, the MB49 and UPPL1541 results indicate that VNB-mediated disruption can induce surface exposure of the pro-phagocytic “eat-me” signal CALR across molecularly distinct bladder cancer models.

Importantly, consensus guidelines indicate that DAMP emission alone is not sufficient to define ICD; functional protection in immunocompetent syngeneic hosts is required.^18^ In agreement with this, vaccination with VNB-killed MB49 cells reduced tumor engraftment and growth after subcutaneous challenge, and splenocytes from VNB-vaccinated mice showed enhanced antitumor activity *ex vivo* (Fig. 4). This protection was also preserved in male mice (Supplementary Fig. S7). In an orthotopic bladder cancer model, VNB vaccination prevented MRI-detectable tumor formation in all vaccinated mice after intravesical challenge (Fig. 6). These protective outcomes were accompanied by early changes in the bladder immune infiltrate. VNB vaccination was associated with a marked increase in bladder-infiltrating CD8⁺ T cells and increased frequencies of CD44⁺CD62L⁺ CD8⁺ T cells (Fig. 7; Supplementary Fig. S9). This phenotype is consistent with a central memory-like population, although its detection at this early time point does not establish the formation of durable immunological memory. Vaccination was also associated with increased frequencies of NK1.1⁺CD62L⁺ cells in the bladder. Given the role of CD62L in lymphoid trafficking and recirculation, this finding may reflect a vaccination-associated shift in the phenotype of the infiltrating NK1.1⁺ compartment.^46^ Alongside these lymphoid changes, we observed increased infiltration of IMs, including the MHC-II⁻ subset, in the bladder 48 h after orthotopic MB49 challenge (Fig. 7 and Supplementary Fig. S8). Domingos-Pereira et al. likewise detected CD11bʰⁱLy6Cʰⁱ monocytic cells in an orthotopic MB49 model at day 5, the earliest time point they examined and reported a marked increase in their abundance by day 9.^47^ The early timing of our observation is consistent with inflammatory monocyte recruitment. However, the MHC-II⁻ subset shares phenotypic markers with monocytic myeloid-derived suppressor cells, so we cannot rule out their presence in the early infiltrate.

Together, our results show that VNB-mediated killing converts MB49 cells into a functionally immunogenic vaccine substrate, linking DAMP emission to protection against tumor challenge in immunocompetent syngeneic hosts and thereby meeting a key functional criterion for ICD.

Whole-tumor antigen strategies are attractive because they provide a broad repertoire of tumor-associated and specific antigens, potentially enabling polyclonal CD8⁺ and CD4⁺ T-cell responses and reducing immune escape through loss of a single antigen.^32,33^ However, antigen breadth alone is insufficient, and the limited clinical efficacy of many whole-tumor-cell vaccines has been linked to weak danger signaling, insufficient APC activation and inefficient cross-priming.^32^ VNB treatment addresses this limitation by combining broad antigen availability with a physical cell-stress modality that generates ICD-associated adjuvanticity. Thus, the DAMP profile observed *in vitro* (Fig. 3; Supplementary Fig. S4) was associated with systemic antitumor activity (Fig. 4), complete orthotopic prophylaxis (Fig. 6), and early recruitment of CD8⁺ T cells and memory-like lymphoid populations to the bladder (Fig. 7).

This strategy is also consistent with recent efforts to improve whole-tumor vaccines by engineering the tumor-cell substrate itself. Genetic engineering has also been explored in bladder cancer: Peng et al. reported that irradiated MB49 cells co-expressing GM-CSF and IL-21 inhibited tumor growth more effectively than vaccines expressing either cytokine alone, whereas unmodified irradiated MB49 cells showed no antitumor effect under their experimental conditions.^48^ Our approach instead uses VNB-mediated photomechanical damage to generate ICD-associated danger signals without cytokine transfection, with protective efficacy evaluated in prophylactic subcutaneous and orthotopic bladder challenge models. Haptenized whole-tumor-cell vaccines based on squaric esters increased antigen recognition, induced immunogenic tumor-cell death, eliminated tumorigenic potential and promoted antigen cross-presentation, achieving strong prophylactic protection in melanoma models.^49^ Glycopolymer-engineered whole-tumor-cell vaccines enhanced interactions with macrophages, promoted M1-like polarization and improved antitumor efficacy.^50^ ConvitVax is another personalized whole-tumor vaccine concept based on autologous tumor-cell homogenate combined with *Mycobacterium bovis* Bacillus Calmette-Guérin (BCG) and low-dose formalin, with preclinical evidence of immune activation and safety in mice.^51^ Compared with these approaches, VNB-mediated cell killing for vaccine manufacturing does not require chemical haptenization, genetic modification, BCG/formalin formulation or extensive membrane engineering. Instead, it uses optical activation of IONPs to inactivate tumor cells, induce DAMP exposure/release and preserve the breadth of the antigen source. Moreover, the optical nature of VNB-mediated killing may offer manufacturing advantages as cell-based products face challenges related to reproducibility, scalability, cost of goods, release testing, preservation and chain-of-custody control.^52^ A pulsed-laser killing step is externally triggered, rapid, spatially controllable and potentially compatible with automated closed or semi-closed workflows. Parameters such as fluence, pulse number, irradiation area, cell density and IONP dose could be standardized, making the process more controllable than manual freeze–thaw cycles or multi-step chemical modifications.

Beyond antitumor efficacy, the tolerability of the vaccine preparation is particularly relevant for whole-cell vaccination strategies, because the administered material contains large amounts of tumor-cell-derived components and, depending on the manufacturing method, may retain residual treatment-associated agents. Previous safety studies of whole-tumor-cell vaccines have therefore combined clinical observation with injection-site assessment, body- and organ-weight measurements, plasma biochemistry and histopathological analysis of major organs. For example, ConvitVax showed only mild, self-limiting local reactions and no treatment-related abnormalities in body weight, clinical chemistry, organ morphology or tissue architecture, supporting this multidimensional approach to vaccine safety evaluation.^51^ In our study, a single administration of VNB-killed MB49 cells was similarly well tolerated over the 7-day observation period, with stable body weight, preserved kidney, liver and spleen histology, unchanged organ weights and no biochemical evidence of hepatic or renal injury. These findings support the short-term safety of both vaccine preparations under the conditions tested, while suggesting a potentially cleaner local tolerability profile for the drug-free VNB-engineered vaccine. Future studies should define the durability of protection, therapeutic activity against established disease, compatibility with intravesical immunotherapies and translation to patient-derived tumor material.

## CONCLUSION

In summary, laser-induced VNBs generated from dextran-coated IONPs provide an externally controllable physical route to convert bladder tumor cells into an immunogenic whole-cell tumor vaccine. Under optimized conditions, VNB treatment induced robust MB49 cell killing together with coordinated CALR exposure and ATP and HMGB1 release, translating into protective antitumor immunity after subcutaneous challenge and prevention of MRI-detectable orthotopic bladder tumor formation in all vaccinated animals. The associated early infiltration of CD8⁺ T cells, memory-like CD8⁺ T-cell populations and inflammatory monocytes further supports the induction of a protective antitumor immune response. A single administration of the VNB-engineered vaccine also showed a favorable short-term systemic safety profile, with no gross injection-site alterations observed in VNB-vaccinated animals. By combining whole-cell-derived tumor material with ICD-associated adjuvanticity, VNB-mediated optical killing offers a drug-free and potentially scalable manufacturing strategy for prophylactic whole-tumor-cell vaccination.

## METHODS

### Iron oxide nanoparticles

Commercially available dextran-coated iron oxide nanoparticles (IONPs, 500 nm; Absolute Mag™ Plain Magnetic Nanoparticles, CD Bioparticles) were used.

### Irradiation set-up

Irradiation was performed using an in-house built pulsed-laser set-up coupled to a THUNDER Leica microscope (Figure S1). A Q-switched, diode-pumped Cobolt Tor™ laser (λ = 532 nm; pulse width 4 ± 1 ns; pulse energy 60 ± 5 µJ) was routed through a tunable 532-nm beam splitter (VA5-532/M, Thorlabs) and relayed to the microscope via a periscope (RS99/M). The beam was delivered through a 10× objective, resulting in an irradiation spot of ∼110 µm diameter at the sample plane. Laser fluence was adjusted by varying the beam-splitter setting and quantified at the sample using a high-speed energy meter equipped with an integrated PE012 pyroelectric sensor (SLT Sensor-und Lasertechnik). The following fluences were used: 0.01, 0.02, 0.16, 0.39, 0.56, and 0.61 J/cm². Vapor nanobubble (VNB) formation was visualized with a DFC9000 CTC camera using a 10× phase-contrast imaging path and a filter cube (ZET532/10×, ZT532-UF3, ET560lp).

### Cell culture

MB49 cells (murine urothelial carcinoma cell line) and UPPL1541 cells (murine bladder cancer cell line with similar expression profiles to luminal-like molecular subtype bladder cancers in humans)^44^ were cultured in Dulbecco’s modified Eagle’s medium (DMEM) containing D-glucose, L-glutamine, and sodium pyruvate. Complete medium was prepared by supplementing DMEM with 10% FBS and penicillin–streptomycin.

### Viability in MB49 cells

To identify the optimal laser fluence for inducing MB49 cell death, cells were seeded at 4.0 × 10⁴ cells/well in glass-bottom 18-well µ-plates (IBIDI) 24 h prior to irradiation. On the day of the experiment, 200 µL IONPs (1:100 dilution from a 10 mg/mL stock in non-supplemented medium) were added per well and incubated for 10, 20, or 30 min. The IONP suspension was aspirated and replaced with complete medium, followed by irradiation at 0.01, 0.02, 0.16, 0.39, 0.56, or 0.61 J/cm².

Cell viability was quantified using the CellTiter-Glo® luminescent viability assay (Promega, Leiden, The Netherlands) according to the manufacturer’s instructions. Briefly, cells were washed with PBS and incubated with 100 µL medium plus 100 µL CellTiter-Glo® reagent per well. Plates were mixed for 10 min on an orbital shaker (120 rpm), and lysates were transferred to white opaque 96-well plates (Costar®) for luminescence acquisition. After 10 min equilibration at room temperature, luminescence was recorded on a Tecan SPARK® multimode plate reader.

For optical visualization of cell killing, cells were stained immediately after irradiation with Ethidium Homodimer-2 (EthD-2; 1 µM; Invitrogen) for 15 min at 37 °C, 5% CO₂. Images were acquired on the THUNDER microscope using a 10× objective and the DFC9000 CTC camera; EthD-2 was excited using a 575-nm light source. Image analysis was performed in FIJI/ImageJ (https://fiji.sc/).

The selected condition for experiments was 20 min incubation and irradiation at a laser fluence of 0.39 J/cm^2^.

MB49 cells were treated as described above and were evaluated at 2, 4 and 6 h after treatment using the CellTiter-Glo luminescent viability assay according to the manufacturer’s instructions to evaluate time-course metabolic viability.

### Annexin V-FITC/PI apoptosis assay

Cells were seeded at 1 × 10^6^ cells per well in 35 mm high glass-bottom µ-Dishes (ibidi) and irradiated as described above. After 2, 4, 6 or 24 h, cells and supernatants were collected and washed twice with PBS. Cells were resuspended in Annexin V binding buffer and stained with Annexin V–FITC (ref. 640945, BioLegend) for 15 min at room temperature in the dark. Propidium iodide (PI) was added to a final concentration of 1 µg/mL. Of note, doxorubicin (DOX) exhibits strong intrinsic fluorescence in the FITC and PI channels. To accurately differentiate true Annexin V and PI staining from DOX background emission, gating thresholds for DOX-treated samples were adjusted using unstained DOX-treated control cells. Samples were analyzed using a four-laser Aurora spectral flow cytometer (Cytek), and data were processed by FlowJo v10.

### DAMPs analysis

*Surface Calreticulin (CALR) Exposure*: CALR exposure was analyzed by spectral flow cytometry at 2, 4, 6 and 24 h after treatment. Cells and their corresponding supernatants were harvested to include both adherent and detached cells, centrifuged at 350 × g for 5 min and resuspended in ice-cold PSA buffer composed of PBS supplemented with 5% FBS and 0.01% sodium azide. Cells were incubated for 15 min at 4 °C to reduce non-specific antibody binding and then stained with Alexa Fluor 405-conjugated anti-CALR antibody (5 µg/mL; Abcam) for 30 min in the dark without permeabilization. After washing, cells were stained with Annexin V–FITC and propidium iodide (PI) to discriminate viable and dying/dead-cell populations. Samples were acquired using a four-laser Aurora spectral flow cytometer (Cytek), and data were analyzed using FlowJo. Cells were first selected according to forward- and side-scatter parameters, followed by singlet discrimination. Viable cells were defined as Annexin V–FITC⁻PI⁻, whereas dying/dead cells were defined as Annexin V–FITC⁺/PI⁺. The frequency of CALR⁺ cells was quantified separately within the viable and dying/dead-cell populations, as shown in Supplementary Fig. S4a.

### ATP release

Cells were seeded in 35 mm high glass-bottom µ-Dishes (Ibidi) at a density of 1 × 10^6^ cells and irradiated as described above. At 2, 4, 6 and 24 h post-irradiation, supernatants were collected and clarified by centrifugation (12,000 × g, 5 min). Extracellular ATP was quantified in white 96-well plates (Costar) using an ATP determination kit according to the manufacturer’s instructions. Luminescence was measured using a Tecan SPARK® multimode reader, and ATP concentrations were determined via a standard calibration curve. Non-irradiated cells served as the negative control, while cells treated with 120 µM doxorubicin were used as a positive control.

### HMGB1 release

Extracellular HMGB1 was quantified using the Lumit® HMGB1 Immunoassay (W6110; Promega) following the manufacturer’s protocol. Supernatants were collected at the indicated time points (2, 4, 6 and 24 h post-irradiation), clarified by centrifugation (≈12,000 × g, 5 min), and stored at 2–10 °C until analysis (≤48 h; no freeze–thaw). Assays were run in white 96-well plates (final volume 125 µL/well). For each well, 80 µL sample (or control) was mixed with 20 µL of 5× antibody mixture (anti-HMGB1 mAb-SmBiT and mAb-LgBiT diluted 1:50 to achieve a 1:250 final antibody dilution) and incubated 60–90 min at room temperature. 25 µL of 5× Detection Reagent B (Detection Substrate B diluted 1:20 in Detection Buffer B) was then added and, after 3–5 min, luminescence was recorded on the Tecan SPARK®.

### Evaluation of VNB-mediated killing and CALR exposure in UPPL1541 cells

To evaluate the reproducibility of VNB-mediated cell killing and CALR exposure in an additional bladder cancer model, UPPL1541 cells were treated using the same experimental conditions described for MB49 cells. Cells were incubated with IONPs at 0.1 mg/mL for 20 min, washed to remove non-associated particles and irradiated at a laser fluence of 0.39 J/cm². Untreated cells, cells incubated with IONPs without irradiation, VNB-killed cells and cells treated with 120 µM doxorubicin were evaluated at 2, 4 and 6 h after treatment. Metabolic viability was quantified using the CellTiter-Glo luminescent viability assay. In parallel, adherent and detached cells were collected together and stained with Annexin V–FITC and PI as described above. Late-apoptotic cells were defined as Annexin V–FITC⁺PI⁺. For analysis of surface CALR exposure, non-permeabilized cells were stained with Alexa Fluor 405-conjugated anti-CALR antibody before Annexin V–FITC/PI staining. CALR positivity was quantified within the dying/dead-cell population. Samples were acquired using a four-laser Aurora spectral flow cytometer and analyzed using FlowJo v10.

### Vaccination-challenge subcutaneous experiments

Animal experiments were conducted in accordance with procedures approved by the Animal Care and Ethics Committee of the Autonomous University of Barcelona (CEEAH) and the Animal Care Committee of the Generalitat de Catalunya. All experiments followed the 3Rs principle to ensure animal welfare. Animals were maintained in quarantine for 5 days to confirm a healthy status, and experiments were carried out in a Biosafety Level 2 facility. Animal procedures were performed in a Class II Biosafety Cabinet due to the handling of tumor cells.

C57BL/6 females and males mice (6–7-week-old, Charles Rivers) were randomized. All procedures were performed under isoflurane anesthesia (1.5-2%) in pure oxygen. After positioned the mice in the prone position and shaving their left flank, 100 µL of culture media containing 1×10⁶ killed MB49 cells was injected into the left flank using a 25G needle. The following groups were included: DOX-killed cells (positive control), VNB-killed cells and vehicle alone (negative control). One week after subcutaneous vaccination, mice were challenged by subcutaneous injection into the right flank with 3×10⁵ viable MB49 cells suspended in 100 µL of DMEM, administered using a 25G needle. Tumor size at the site of subcutaneous challenge was measured twice weekly using a caliper. Two perpendicular tumor diameters (length and width) were recorded, and tumor size was calculated as length × width and expressed in mm². Animals were housed in groups of up to five-six per cage, with free access to food and water. Body weight, appearance, and behavior were regularly monitored. Clinical signs were assessed daily to ensure that the animals did not exhibit signs of stress or pain, and animals were euthanized when it was required. At the end of the experiments, animals were euthanized, and spleens were removed under aseptic conditions.

### Splenocyte-mediated antitumor activity

Spleens were mechanically dissociated using forceps. The resulting splenocyte suspensions were homogenized and diluted in complete RPMI medium and then seeded into 96-well tissue culture plates (Thermo Scientific) containing 1×10^4^ MB49 cells at a splenocyte-to-tumor cell ratio of 25:1. After 48 h of incubation at 37°C in a humidified atmosphere with 5% CO₂, wells were thoroughly washed with warm PBS to remove non-adherent splenocytes. The proliferation of the remaining adherent cell population, which includes proliferating MB49 cells and probably a small proportion of adherent monocytes/macrophages and other myeloid cells present in the splenocyte preparation, was assessed using the MTT assay.

### Safety evaluation

Female C57BL/6J mice (6–7 weeks old; Janvier Labs) were housed in the animal research facilities of IQAC-CSIC. All animal procedures were evaluated and approved by the Animal Experimentation Ethics Committee of the Generalitat de Catalunya (protocol OH-2107/2026). Animals were randomized into three experimental groups (n = 8 per group) and received a single subcutaneous injection of 1 × 10⁶ VNB-killed MB49 cells, DOX-killed MB49 cells or an equivalent volume of DMEM as vehicle control. Vaccine preparations and administration were performed as described for the subcutaneous vaccination–challenge experiments. Body weight was recorded before administration and on days 2, 5 and 7, and the general condition of the animals and the appearance of the injection site were monitored throughout the 7-day observation period. At the day-7 endpoint, mice were euthanized and blood was collected by cardiac puncture. Plasma samples were analyzed at the Servei de Bioquímica Clínica Veterinària, Facultat de Veterinària, Universitat Autònoma de Barcelona. Total protein, albumin, creatinine, glucose, aspartate aminotransferase (AST), alanine aminotransferase (ALT), total bilirubin, urea and cholesterol were quantified to assess hepatic and renal function and general metabolic status. The liver, kidneys and spleen were collected, macroscopically examined and weighed at IQAC-CSIC. Representative samples of these organs were preserved for histopathological evaluation. Tissue processing, paraffin embedding, sectioning and hematoxylin and eosin staining were performed by the Histopathology Facility of the Institute for Research in Biomedicine Barcelona (IRB Barcelona). Sections were evaluated for preservation of tissue architecture and for evidence of treatment-associated inflammation, cellular degeneration, necrosis or other pathological alterations.

### Vaccination-challenge orthotopic experiments

C57BL/6 female mice (6–7-week-old, Charles Rivers) mice were first randomized and were subcutaneously injected with treated dead MB49 cells as described above. One week after subcutaneous vaccination, mice were challenged by intravesical instillation of 1×10⁵ viable MB49 cells suspended in 100 µL of DMEM. Tumor cells were delivered into the bladder via a 24-gauge catheter (Becton Dickinson), as previously described.^53^ All procedures were performed under isoflurane anesthesia (1.5-2%) in pure oxygen and eyes protected with liquid ophthalmic gel. Animals were housed in groups of up to five-six per cage, with free access to food and water. Body weight, appearance, and behavior were regularly monitored. Clinical signs were assessed daily to ensure that the animals did not exhibit signs of stress or pain, and animals were euthanized when it was required.

Two sets of experiments were performed. In the first set, animals were euthanized two days after intravesical MB49 tumor challenge, and bladders were collected under aseptic conditions to evaluate immune cell infiltration. In the second set, animals were evaluated at days 21-24 after tumor induction to assess tumor volume by magnetic resonance imaging (MRI) and were euthanized at day 30 after challenge.

### Intravesical tumor size

MRI studies were conducted 21–24 days after intravesical tumor challenge, at the joint nuclear magnetic resonance facility of the Universitat Autònoma de Barcelona and Centro de Investigación Biomédica en Red—Bioingeniería, Biomateriales y Nanomedicina (CIBER-BBN) (Cerdanyola del Vallès, Spain), Unit 25 of NANBIOSIS. Experiments were performed on a 7T Bruker Biospec 70/30 USR preclinical scanner using ParaVision 5.1 (Bruker BioSpin GmbH, Ettlingen, Germany) equipped with a BGA-12 gradient insert of 440 mT m^−1^, and a 72 mm inner diameter volume resonator (T10720) for radiofrequency transmission and a rat brain surface coil (T11205) for RF reception (both operating at 300 MHz). Animals were anaesthetized with isoflurane (4% for induction and 1.5% for maintenance in O2 at 1 L/min) and placed on an MR-compatible cradle. Their respiration rates were constantly monitored with a pressure probe and kept between 60 and 80 breaths/min (SA Instruments, Inc., New York, USA). Mouse body temperature was monitored using a rectal probe and maintained at 37 ± 1 °C with a heated recirculating water system incorporated in the animal bed. After acquiring reference images, high-resolution T2-weighted (T2w) images were acquired in axial planes using a rapid acquisition with relaxation enhancement (RARE) sequence with the following parameters: RARE factor, 12; effective echo time (TEeff), 40 ms; repetition time (TR), 3600 ms; field of view (FOV), 16×16 mm^2^; matrix size (MTX), 160×160 (in-plane resolution, 0.1×0.1 mm^2^/pixel); number of slices (NS), 24; slice thickness (ST), 0.5 mm (contiguous slices with no gap, acquired in interleaved mode); number of averages (NA) 16; acquisition time (AT), 12 min 29 s. Furthermore, a spin-echo-based diffusion-weighted (DW) imaging sequence was used to enhance tumour visualization using the following parameters: TE, 20 ms; TR, 2550 ms; FOV, 16×16 mm2; MTX, 160×160; 10 contiguous 1 mm slices; NA, 2. For each slice, one A0 image (basal image with b = 0 s mm^−2^) and one DW image at b = 650 s mm^−2^ were acquired using diffusion gradients in the (1, 0, 0) direction with a gradient duration δ, 3.5 ms; and a gradient separation Δ , 9 ms; AT, 27 min 12 s. Tumors were manually delineated to determine their volume.

### Detection of infiltrated immune cell subsets into the bladders

The immune infiltrate within the bladder was characterized by flow cytometry as previously reported.^53^ In brief, bladder tissues were mechanically dissociated using a scalpel and subsequently enzymatically digested in RPMI medium supplemented with 5% fetal bovine serum (FBS), 0.5 mg/mL collagenase II (Sigma, Spain), and 1 U/mL DNase I. Digestion was performed at 37°C under continuous agitation in two to three consecutive 30-minute cycles. The resulting cell suspension was passed through a 40-µm cell strainer (Becton Dickinson) to remove debris and obtain a single-cell suspension, followed by centrifugation for downstream staining procedures. Cells were incubated with fluorochrome-conjugated monoclonal antibodies (Table S1). Lymphocytes were identified based on forward and side scatter morphological parameters, and non-viable cells were excluded using the Live/Dead Fixable Aqua Dead Cell Stain kit (Invitrogen). Data acquisition was performed on a Fortessa flow cytometer (Becton Dickinson), and analysis was conducted using FlowJo software (version 9.8; Tree Star, Portland, OR, USA).

### Statistical analysis

All graphs were made using Prism 10.1.1 (GraphPad Software). Data analysis and statistical significance were performed using Prism 10.1.1, GraphPad Software, where *p<0.05, **p<0.01, ***p<0.001, and ****p<0.0001.

## ACKNOWLEDGMENTS

This research was funded by grants PID2021-122331OB-I00 (Spanish Ministry of Science) and 2021SGR-0092 (AGAUR) from E.J., HR21-00208 (Fundación “La Caixa”) from E.J. and S.S, and PID2024-158142OA-I00 from J.C.F. E.J. is a Serra-Húnter fellow. G.C. acknowledges support from Grant PTA2024-024509-I funded by MICIU/AEI/10.13039/501100011033 and by the ESF+. J.C.F. gratefully acknowledges the financial support from the Spanish Ministry of Science through the Ramón y Cajal Grant No. RYC2023-043053-I. J.C.F. (2021BP-00079), N.S.G. (2023FI-100137), N.T. (2024FI-100310) and A.G.C. (2023DI-00122) acknowledge AGAUR (Generalitat de Catalunya) for their support.

## AUTHOR CONTRIBUTIONS

J.C.F. conceived and led the study, coordinated the *in vitro* and technological work, supervised the development of the VNB platform, contributed to the conceptualization of the *in vitro* and *in vivo* experimental strategy, acquired funding, provided overall guidance, wrote the manuscript and performed the final revision. E.J.G. co-led the study, coordinated the *in vivo* and immunological studies, contributed to the conceptualization of the *in vivo* strategy together with N.S.G., A.G.C. and J.C.F., supervised the animal experiments, acquired funding, wrote the manuscript and contributed to manuscript revision. S.S. provided senior scientific guidance, acquired funding and contributed to manuscript revision. N.S.G. performed the *in vivo* experiments under the supervision of E.J.G. and contributed to manuscript writing. A.G.C. performed the *in vitro* experiments under the supervision of J.C.F. and contributed to manuscript writing. N.T. contributed to the *in vivo* experiments together with N.S.G. and E.J.G. A.G.H., J.R., K.B. and A.F.V. contributed to the design and discussion of the *in vitro* protocols together with A.G.C. and J.C.F., and provided scientific guidance. D.P. and K.B. contributed to the design and development of the optical set-up together with J.C.F. C.C. and J.P. contributed to the flow-cytometry analysis of digested bladder cell populations. G.C. and I.A. contributed to the safety evaluation of VNB-killed cells as whole-tumor-cell vaccines. All authors discussed the results and reviewed the manuscript.

## ADDITIONAL INFORMATION

### Declaration of AI-assisted language editing

During manuscript preparation, the authors used ChatGPT (OpenAI) to assist with English language editing and sentence restructuring. All scientific content, interpretations, references and final wording were reviewed and verified by the authors, who take full responsibility for the submitted work.

## Supplementary information

The online version contains supplementary material, including the optical layout of the laser irradiation system, optimization of doxorubicin- and vapor-nanobubble-mediated MB49 cell killing, representative flow-cytometry gating strategies for cell-death, calreticulin and immune-infiltrate analyses, calibration curves for ATP and HMGB1 quantification, additional subcutaneous vaccination–challenge data in male mice, *in vivo* safety assessment, and the antibody panels used for immune profiling.

## REFERENCES

1 Sung H, Filho AM, Laversanne M, Ferlay J, Siegel RL, Soerjomataram I et al. Global cancer statistics 2024: GLOBOCAN estimates of incidence and mortality worldwide for 34 cancers in 186 countries. CA Cancer J Clin 2026; 76. doi:10.3322/caac.70090.

2 Chang SS, Boorjian SA, Chou R, Clark PE, Daneshmand S, Konety BR et al. Diagnosis and Treatment of Non-Muscle Invasive Bladder Cancer: AUA/SUO Guideline. Journal of Urology 2016; 196: 1021–1029.

3 Pettenati C, Ingersoll MA. Mechanisms of BCG immunotherapy and its outlook for bladder cancer. Nat Rev Urol 2018; 15: 615–625.

4 Ślusarczyk A, Scilipoti P, Contieri R, Krajewski W, Claps F, Subiela JD et al. Refining BCG-failure classifications in non-muscle-invasive bladder cancer. BJU Int 2026; 137: 1076–1087.

5 Saboya L, Buosi K, Silva T, Candido E, Morari J, Velloso LA et al. Intradermal priming to intravesical Bacillus Calmette-Guérin in non-muscle invasive bladder cancer: A translational research and phase I clinical trial. Oncol Res 2025; 33: 1495–1503.

6 Buisan O, Servian P, Pedreño-Lopez S, Pagès J, Urrea V, Martínez R et al. A Novel Heterologous Prime-boost Strategy Using RUTI Vaccine to Improve the Bacillus Calmette-Guérin Response in Non–muscle-invasive Bladder Cancer Patients: Phase 1 RUTIVAC-1 Trial. Eur Urol 2026; 89: 114–120.

7 Niwa N, Kikuchi E, Matsumoto K, Kosaka T, Mizuno R, Oya M. Purified Protein Derivative Skin test prior to bacillus Calmette-Guérin Therapy May have Therapeutic Impact in Patients with Nonmuscle Invasive Bladder Cancer. Journal of Urology 2018; 199: 1446–1451.

8 Ribas A, Wolchok JD. Cancer immunotherapy using checkpoint blockade. Science (1979) 2018; 359: 1350–1355.

9 Li X, Lovell JF, Yoon J, Chen X. Clinical development and potential of photothermal and photodynamic therapies for cancer. Nat Rev Clin Oncol 2020; 17: 657–674.

10 Yun WS, Park JH, Lim DK, Ahn CH, Sun IC, Kim K. How Did Conventional Nanoparticle-Mediated Photothermal Therapy Become “Hot” in Combination with Cancer Immunotherapy? Cancers (Basel) 2022; 14: 1–20.

11 Zhao L, Zhang X, Wang X, Guan X, Zhang W, Ma J. Recent advances in selective photothermal therapy of tumor. J Nanobiotechnology 2021; 19: 1–15.

12 Han HS, Choi KY. Advances in nanomaterial-mediated photothermal cancer therapies: Toward clinical applications. Biomedicines 2021; 9: 1–15.

13 Sweeney EE, Cano-Mejia J, Fernandes R. Photothermal Therapy Generates a Thermal Window of Immunogenic Cell Death in Neuroblastoma. Small 2018; 14: 1–8.

14 Zhang Y, Zhan X, Xiong J, Peng S, Huang W, Joshi R et al. Temperature-dependent cell death patterns induced by functionalized gold nanoparticle photothermal therapy in melanoma cells. Sci Rep 2018; 8: 1–9.

15 Deng X, Shao Z, Zhao Y. Solutions to the Drawbacks of Photothermal and Photodynamic Cancer Therapy. Advanced Science 2021; 8: 1–16.

16 Kroemer G, Galassi C, Zitvogel L, Galluzzi L. Immunogenic cell stress and death. Nat Immunol 2022; 23: 487–500.

17 Rodrigues MC, Morais JAV, Ganassin R, Oliveira GRT, Costa FC, Morais AAC et al. An Overview on Immunogenic Cell Death in Cancer Biology and Therapy. Pharmaceutics 2022; 14: 1–17.

18 Galluzzi L, Vitale I, Warren S, Adjemian S, Agostinis P, Martinez AB et al. Consensus guidelines for the definition, detection and interpretation of immunogenic cell death. J Immunother Cancer 2020; 8: 1–22.

19 Krysko D V., Garg AD, Kaczmarek A, Krysko O, Agostinis P, Vandenabeele P. Immunogenic cell death and DAMPs in cancer therapy. Nat Rev Cancer 2012; 12: 860–875.

20 Fucikova J, Kepp O, Kasikova L, Petroni G, Yamazaki T, Liu P et al. Detection of immunogenic cell death and its relevance for cancer therapy. Cell Death Dis 2020; 11. doi:10.1038/s41419-020-03221-2.

21 Nguyen HT, Tran KK, Sun B, Shen H. Activation of inflammasomes by tumor cell death mediated by gold nanoshells. Biomaterials 2012; 33: 2197–2205.

22 Bear AS, Kennedy LC, Young JK, Perna SK, Mattos Almeida JP, Lin AY et al. Elimination of Metastatic Melanoma Using Gold Nanoshell-Enabled Photothermal Therapy and Adoptive T Cell Transfer. PLoS One 2013; 8. doi:10.1371/journal.pone.0069073.

23 Yang W, Guo W, Le W, Lv G, Zhang F, Shi L et al. Albumin-Bioinspired Gd:CuS Nanotheranostic Agent for in Vivo Photoacoustic/Magnetic Resonance Imaging-Guided Tumor-Targeted Photothermal Therapy. ACS Nano 2016; 10: 10245–10257.

24 Tian Y, Zhang J, Tang S, Zhou L, Yang W. Polypyrrole Composite Nanoparticles with Morphology-Dependent Photothermal Effect and Immunological Responses. Small 2016; 12: 721–726.

25 Paholak HJ, Stevers NO, Chen H, Burnett JP, He M, Korkaya H et al. Elimination of epithelial-like and mesenchymal-like breast cancer stem cells to inhibit metastasis following nanoparticle-mediated photothermal therapy. Biomaterials 2016; 104: 145–157.

26 Yu G-T, Rao L, Wu H, Yang L-L, Bu L-L, Deng W-W et al. Myeloid-Derived Suppressor Cell Membrane-Coated Magnetic Nanoparticles for Cancer Theranostics by Inducing Macrophage Polarization and Synergizing Immunogenic Cell Death. Adv Funct Mater 2018; 28: 1801389.

27 Fraire JC, Houthaeve G, Liu J, Raes L, Vermeulen L, Stremersch S et al. Vapor nanobubble is the more reliable photothermal mechanism for inducing endosomal escape of siRNA without disturbing cell homeostasis. Journal of Controlled Release 2020; 319: 262–275.

28 Xiong R, Samal SK, Demeester J, Skirtach AG, De Smedt SC, Braeckmans K. Laser-assisted photoporation: fundamentals, technological advances and applications. Adv Phys X 2016; 1: 596–620.

29 Ramon J, Xiong R, De Smedt SC, Raemdonck K, Braeckmans K. Vapor nanobubble-mediated photoporation constitutes a versatile intracellular delivery technology. Curr Opin Colloid Interface Sci 2021; 54: 101453.

30 Lukianova-Hleb EY, Kim YS, Belatsarkouski I, Gillenwater AM, O’Neill BE, Lapotko DO. Intraoperative diagnostics and elimination of residual microtumours with plasmonic nanobubbles. Nat Nanotechnol 2016; 11: 525–532.

31 Sauvage F, Nguyen VP, Li Y, Harizaj A, Sebag J, Roels D et al. Laser-induced nanobubbles safely ablate vitreous opacities in vivo. Nat Nanotechnol 2022. doi:10.1038/s41565-022-01086-4.

32 Pérez-Baños A, Gleisner MA, Flores I, Pereda C, Navarrete M, Araya JP et al. Whole tumour cell-based vaccines: tuning the instruments to orchestrate an optimal antitumour immune response. Br. J. Cancer. 2023; 129: 572–585.

33 Diao L, Liu M. Rethinking Antigen Source: Cancer Vaccines Based on Whole Tumor Cell/tissue Lysate or Whole Tumor Cell. Advanced Science. 2023; 10. doi:10.1002/advs.202300121.

34 Ramon J, Engelen Y, De Keersmaecker H, Goemaere I, Punj D, Mejía Morales J et al. Laser-induced vapor nanobubbles for B16-F10 melanoma cell killing and intracellular delivery of chemotherapeutics. Journal of Controlled Release 2024; 365: 1019–1036.

35 Stueber DD, Villanova J, Aponte I, Xiao Z, Colvin VL. Magnetic nanoparticles in biology and medicine: Past, present, and future trends. Pharmaceutics. 2021; 13. doi:10.3390/pharmaceutics13070943.

36 Yang X, Kubican SE, Yi Z, Tong S. Advances in magnetic nanoparticles for molecular medicine. Chemical Communications 2025; 61: 3093–3108.

37 Díaz-Riascos Z V., Llaguno-Munive M, Lafuente-Gómez N, Luengo Y, Holmes S, Volatron J et al. Preclinical Development of Magnetic Nanoparticles for Hyperthermia Treatment of Pancreatic Cancer. ACS Appl Mater Interfaces 2025; 17: 2924–2939.

38 Shaterabadi Z, Nabiyouni G, Soleymani M. High impact of in situ dextran coating on biocompatibility, stability and magnetic properties of iron oxide nanoparticles. Materials Science and Engineering C 2017; 75: 947–956.

39 Shi C, Pamer EG. Monocyte recruitment during infection and inflammation. Nat. Rev. Immunol. 2011; 11: 762–774.

40 Qian BZ, Li J, Zhang H, Kitamura T, Zhang J, Campion LR et al. CCL2 recruits inflammatory monocytes to facilitate breast-tumour metastasis. Nature 2011; 475: 222–225.

41 Xiong R, Sauvage F, Fraire JC, Huang C, De Smedt SC, Braeckmans K. Photothermal Nanomaterial-Mediated Photoporation. Acc Chem Res 2023; 56: 631–643.

42 Stewart MP, Langer R, Jensen KF. Intracellular Delivery by Membrane Disruption: Mechanisms, Strategies, and Concepts. Chem Rev 2018; 118: 7409–7531.

43 Garg AD, Krysko D V., Verfaillie T, Kaczmarek A, Ferreira GB, Marysael T et al. A novel pathway combining calreticulin exposure and ATP secretion in immunogenic cancer cell death. EMBO Journal 2012; 31: 1062–1079.

44 Saito R, Smith CC, Utsumi T, Bixby LM, Kardos J, Wobker SE et al. Molecular subtype-specific immunocompetent models of high-grade urothelial carcinoma reveal differential neoantigen expression and response to immunotherapy. Cancer Res 2018; 78: 3954–3968.

45 Xue X, Qu H, Bo R, Zhang D, Zhu Z, Xiang B et al. A transformable nanoplatform with multiple therapeutic and immunostimulatory properties for treatment of advanced cancers. Biomaterials 2023; 299. doi:10.1016/j.biomaterials.2023.122145.

46 Ran G he, Lin Y qing, Tian L, Zhang T, Yan D mei, Yu J hua et al. Natural killer cell homing and trafficking in tissues and tumors: from biology to application. Signal Transduct. Target. Ther. 2022; 7. doi:10.1038/s41392-022-01058-z.

47 Domingos-Pereira S, Sathiyanadan K, Polak L, Haefliger JA, Schmittnaegel M, Ries CH et al. Tumor-Microenvironment Characterization of the MB49 Non-Muscle-Invasive Bladder-Cancer Orthotopic Model towards New Therapeutic Strategies. Int J Mol Sci 2023; 24. doi:10.3390/ijms24010123.

48 Peng J, Ye L, Li T, Zhu Q, Guo J, Xiao K et al. Irradiated Bladder Cancer Cells Expressing both GM-CSF and IL-21 versus Either GM-CSF or IL-21 Alone as Tumor Vaccine in a Mouse Xenograft Model. Biomed Res Int 2019; 2019. doi:10.1155/2019/8262989.

49 Chen Y, Guo S, Tan BSN, Zhao Y, Cheng Y, Zhu W et al. Haptenized whole tumor cell vaccines for cancer immunotherapy. Materials Today 2026; 94. doi:10.1016/j.mattod.2026.103228.

50 Yang H, Feng R, Heng X, Shan F, Wang Y, Yao L et al. Enhanced Whole Tumor Cell-Based Vaccines by a RAFT and Protein Fusion Strategy for Tumor Immunotherapy. Biomacromolecules 2025; 26: 2690–2699.

51 Duarte C MA, Carballo O JM, De Gouveia YM, García A, Ruiz D, Gledhill T et al. Toxicity evaluation of ConvitVax breast cancer immunotherapy. Sci Rep 2021; 11. doi:10.1038/s41598-021-91995-6.

52 Aijaz A, Li M, Smith D, Khong D, Leblon C, Fenton OS et al. Biomanufacturing for clinically advanced cell therapies. Nat. Biomed. Eng. 2018; 2: 362–376.

53 Guallar-Garrido S, Campo-Pérez V, Pérez-Trujillo M, Cabrera C, Senserrich J, Sánchez-Chardi A et al. Mycobacterial surface characters remodeled by growth conditions drive different tumor-infiltrating cells and systemic IFN-γ/IL-17 release in bladder cancer treatment. Oncoimmunology 2022; 11. doi:10.1080/2162402X.2022.2051845.

