## Supporting information for "Photomechanical engineering of immunogenic whole-cell vaccines by laser-induced vapor nanobubbles for bladder cancer prophylaxis"

**Figure S1.**

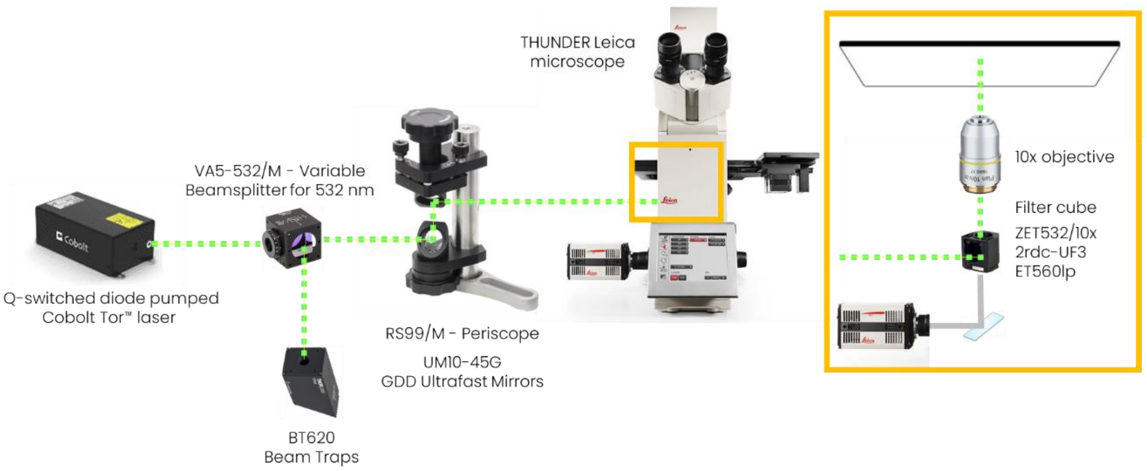


**Fig. S1 | Optical layout for laser-induced vapor nanobubble generation.** Schematic of the pulsed-laser irradiation setup coupled to a THUNDER Leica microscope. A Q-switched diode-pumped Cobolt Tor laser operating at 532 nm was directed through a variable 532-nm beam splitter, relayed through a periscope and ultrafast mirrors, and focused onto the sample through a 10× objective. Laser fluence was adjusted using the variable beam splitter. VNB formation was visualized through the phase-contrast imaging path using a Leica DFC9000 CTC camera and the indicated filter-cube configuration.

**Figure S2.**

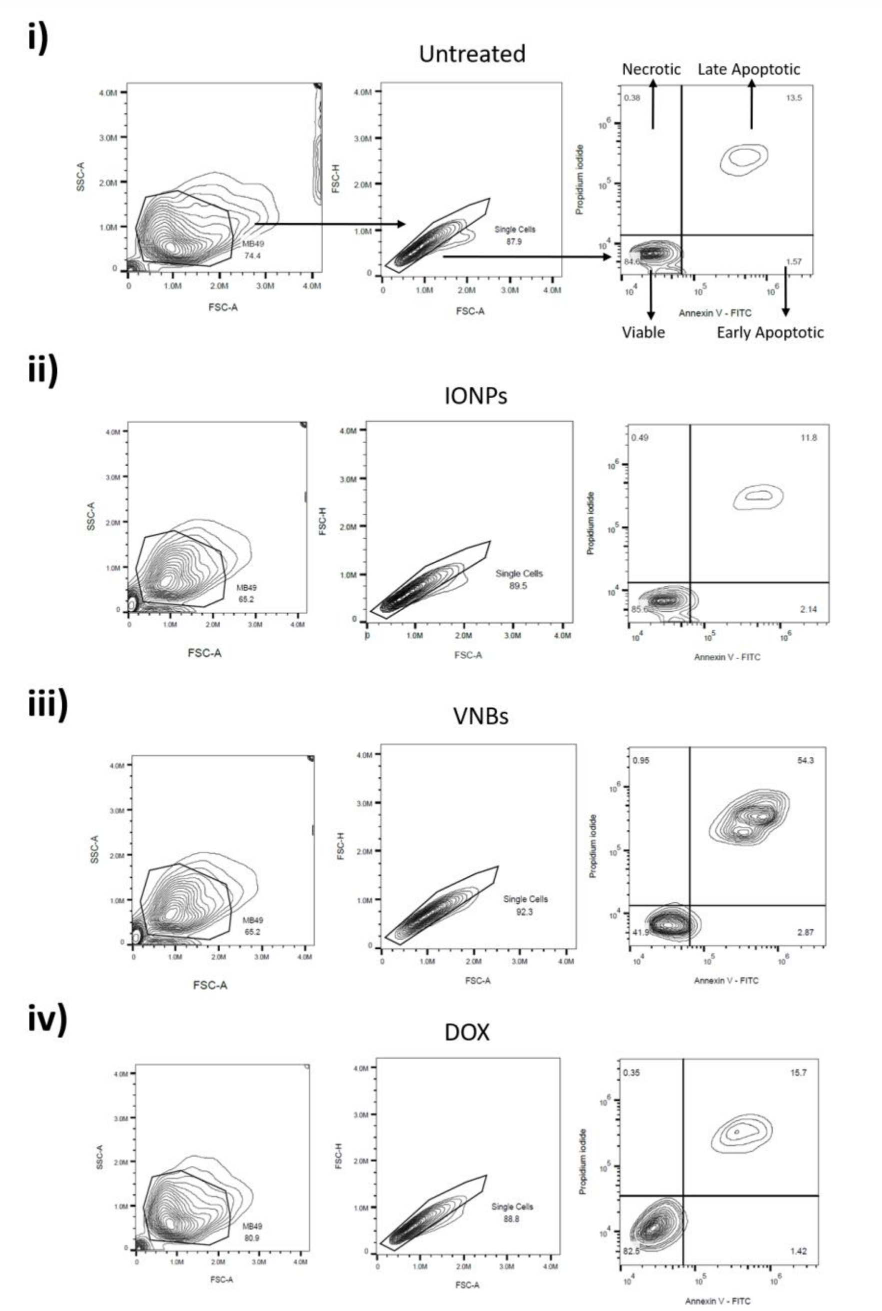


**Fig. S2 | Representative Annexin V and propidium iodide flow-cytometry gating for MB49 cell-death analysis.** Representative gating strategy used to quantify viable, early apoptotic, late apoptotic and necrotic MB49 cells after treatment. Cells were first gated by forward and side scatter, followed by singlet selection. Annexin V–FITC and propidium iodide staining was then used to distinguish viable, early apoptotic, late apoptotic and necrotic cells. Representative plots are shown for (i) non-treated control cells, (ii) IONP-treated cells, (iii) VNB-treated cells and (iv) DOX-treated cells.

**Figure S3.**

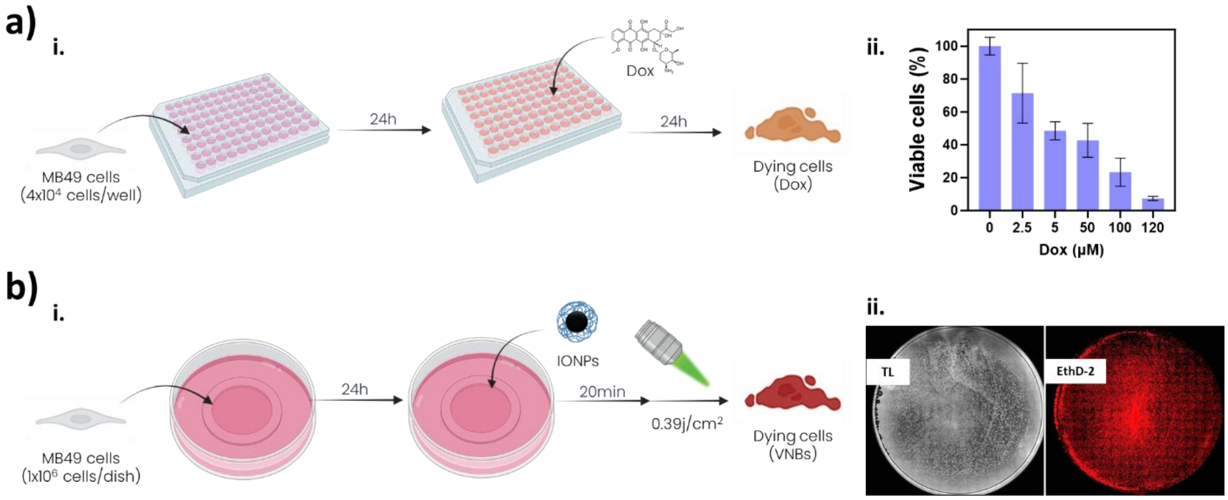


**Fig. S3 | Optimization of doxorubicin- and vapor nanobubble-mediated MB49 cell killing.** a, (i) Doxorubicin killing workflow. (ii) MB49 cells were seeded in 96-well plates, treated with doxorubicin for 24 h and analyzed 24 h later to determine the concentration required to generate dying tumor cells for vaccination studies. Data are shown as mean ± s.d.; n = 3. b, (i) VNB-mediated killing workflow. (ii) MB49 cells were seeded in glass-bottom dishes, incubated with IONPs and irradiated with a pulsed laser to generate VNBs. Representative transmitted-light (TL) and EthD-positive fluorescence (EthD-2) images show widespread cell death after irradiation.

Figure S4.

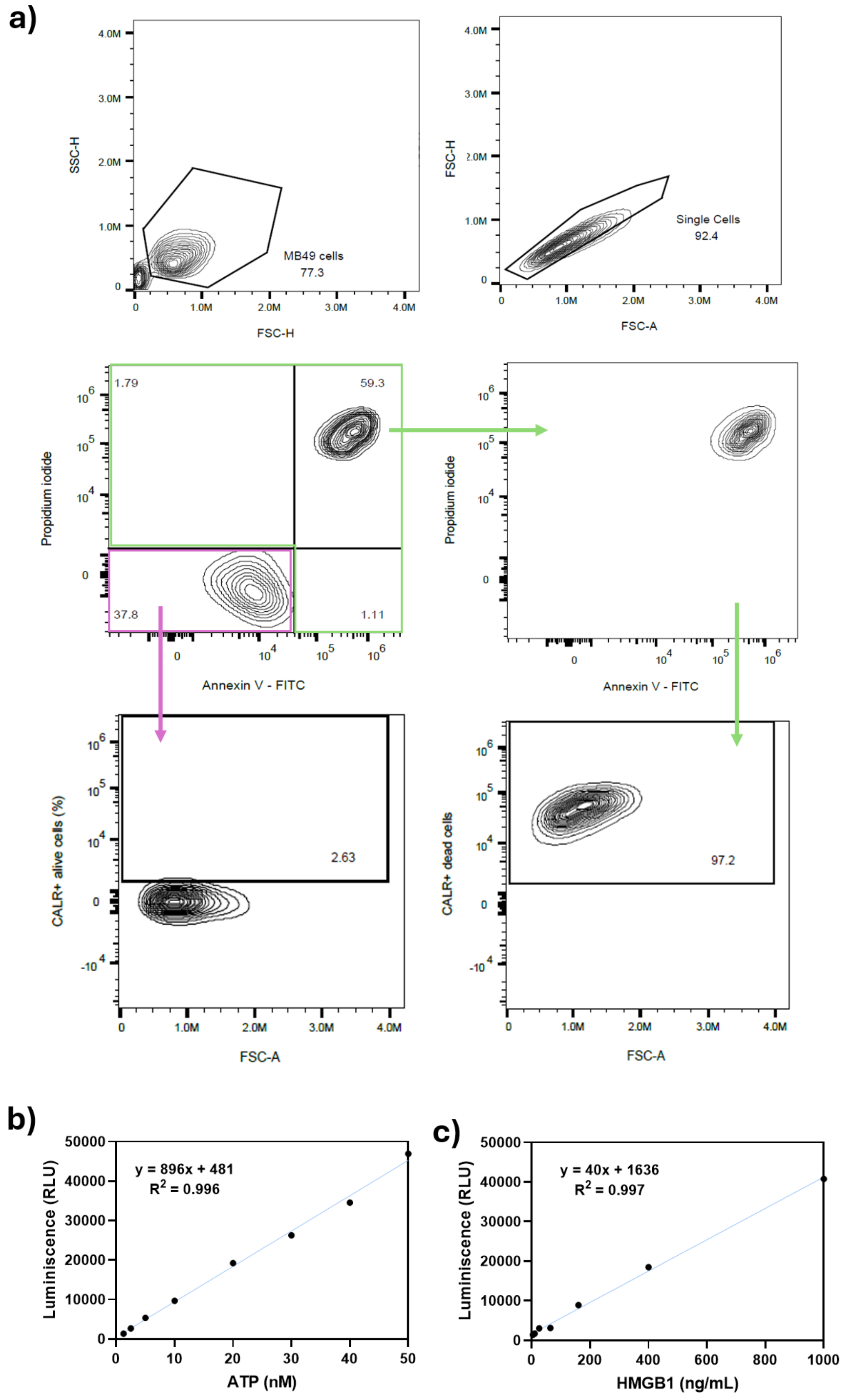


**Fig. S4 | Flow-cytometry gating strategy for CALR analysis and calibration curves for extracellular ATP and HMGB1 release quantification.** a, Representative flow-cytometry gating strategy used to quantify surface CALR exposure after treatment. Cells were first selected according to forward- and side-scatter parameters, followed by singlet discrimination. Live and dead-cell populations were defined using Annexin V–FITC and propidium iodide (PI) staining.

Dead cells were identified using combined Annexin V–FITC$^{+}$/PI$^{+}$gates, and the frequency of CALR$^{+}$ cells was quantified within the corresponding cell populations. Representative plots for the indicated experimental groups are shown. b, ATP standard calibration curve used to quantify extracellular ATP levels in cell-culture supernatants. Luminescence values were plotted against ATP concentration and fitted to the indicated regression model. c, HMGB1 standard calibration curve used to quantify extracellular HMGB1 levels in cell-culture supernatants using the Lumit immunoassay. Luminescence values were plotted against HMGB1 concentration and fitted to the indicated regression model.

**Figure S5.**

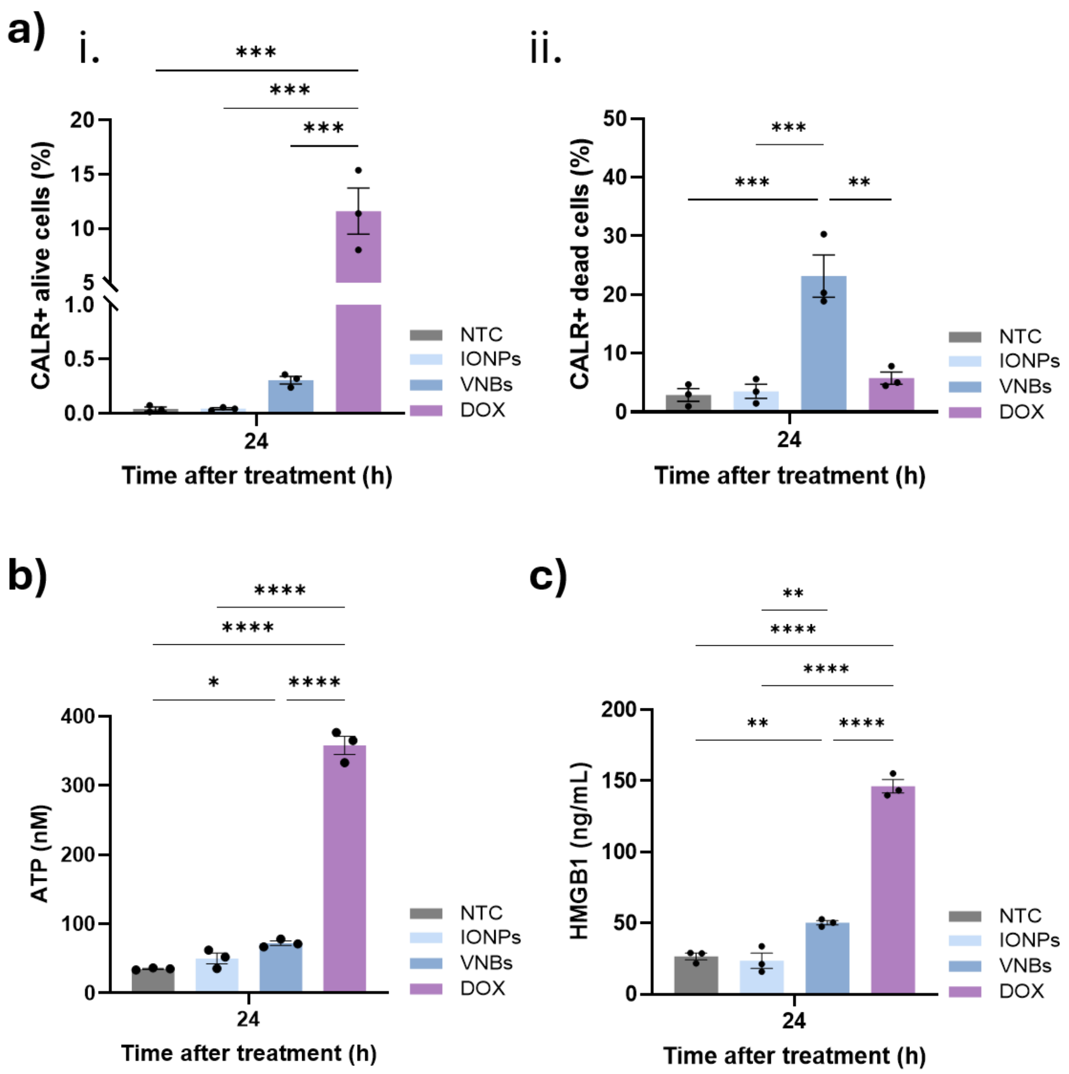


**Fig. S5 | Extended 24-h profiling of ICD-associated DAMP release following VNB-mediated MB49 cell death, with doxorubicin validation as a positive control.** a, Quantification of CALR-positive (i) viable cells and (ii) dying/dead cells 24 h after treatment. b, ATP release measured in cell culture supernatants 24 h after treatment. c, Extracellular HMGB1 release measured in culture supernatants 24 h after treatment. Experimental groups include untreated control cells (NTC), IONP-treated cells (IONPs), VNB-killed cells (VNBs) and doxorubicin-treated cells (DOX). Data are shown as mean ± SEM.; individual points indicate independent biological replicates; n = 3. Statistical significance was determined by one-way ANOVA with Tukey's multiple-comparisons test; *$p < 0.05$, **$p < 0.01$, ***$p < 0.001$ and ****$p < 0.0001$.

**Figure S6.**

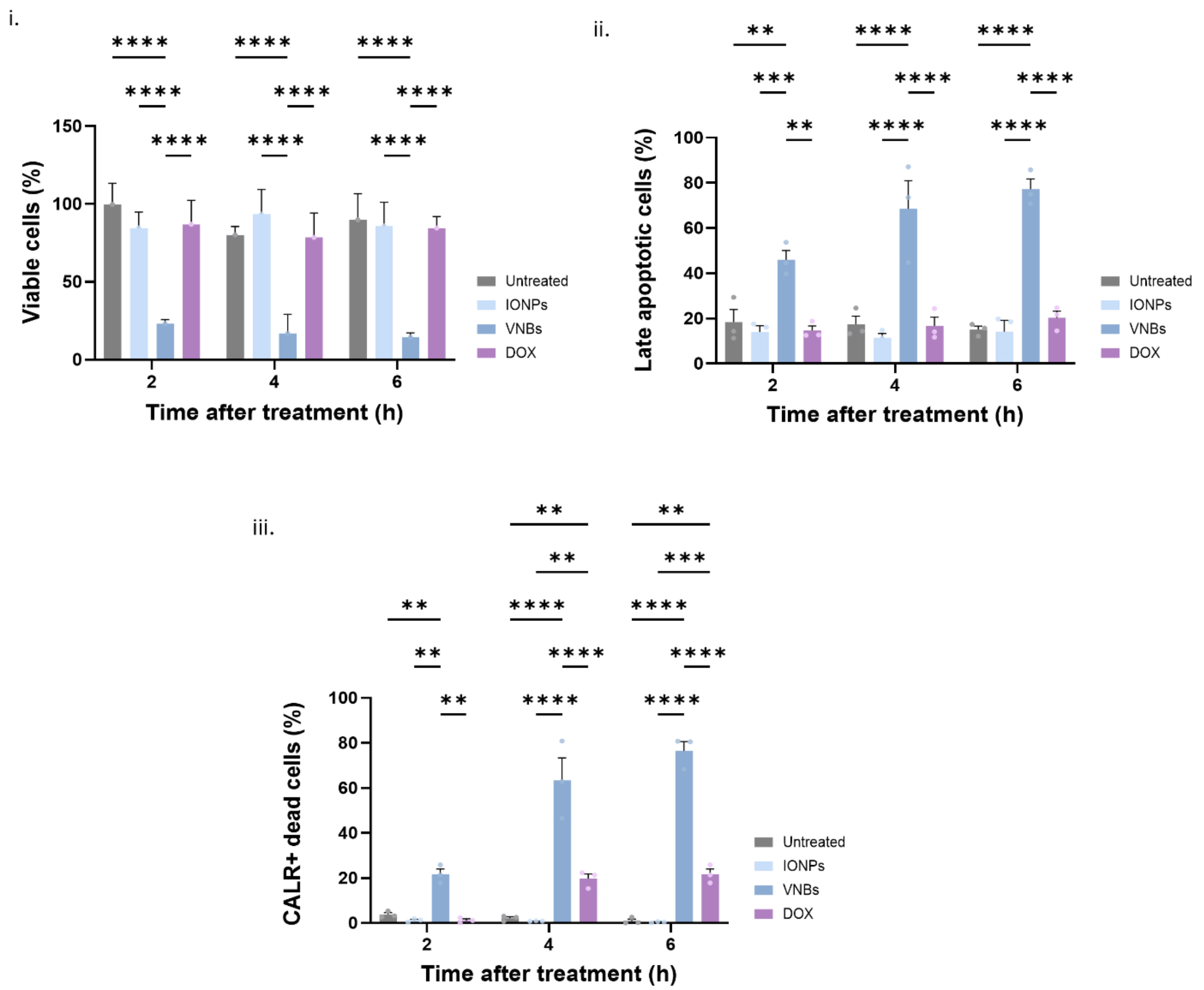


**Fig. S6 | Time-course evaluation of cell killing and CALR exposure in UPPL1541 cells.** (i) Cell viability determined by CellTiter-Glo metabolic viability assay. (ii) Quantification of late apoptotic cells (Annexin V–FITC$^+$/PI$^+$) by flow cytometry. (iii) Quantification of CALR-positive dying/dead cells. Experimental groups include untreated control cells (NTC), IONP-treated cells (IONPs), VNB-killed cells (VNBs) and doxorubicin-treated cells (DOX). Doxorubicin was used as a positive control for immunogenic cell death. Data are shown as mean ± SEM; individual points indicate independent biological replicates; n = 3. Statistical significance was determined by two-way ANOVA followed by Tukey's multiple-comparisons test.*$p < 0.05$, **$p < 0.01$, ***$p < 0.001$ and ****$p < 0.0001$.

**Figure S7.**

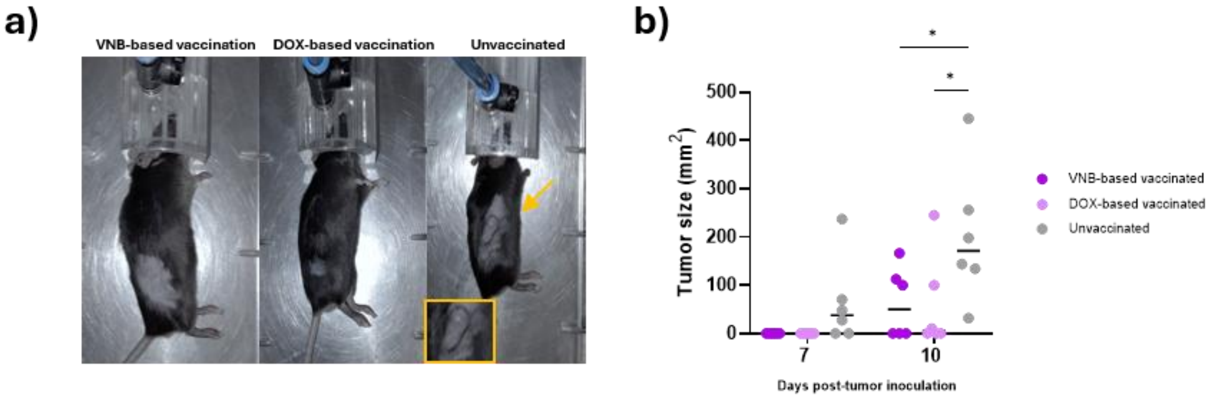


**Fig. S7 | Representative subcutaneous tumors and tumor-size quantification after vaccination–challenge in male mice.** a, Representative images of VNB-based vaccinated, DOX-based vaccinated and unvaccinated male mice after subcutaneous MB49 challenge. Arrows and magnified insets indicate visible tumor lesions. b, Tumor size at the indicated time points after subcutaneous MB49 challenge in VNB-based vaccinated, DOX-based vaccinated and unvaccinated mice. Individual points indicate individual mice and horizontal lines indicate the mean; n = 6 mice per group. Statistical significance was determined by two-way repeated-measures ANOVA followed by Tukey's multiple-comparisons test. *$p < 0.05$, **$p < 0.01$, ***$p < 0.001$ and ****$p < 0.0001$.

**Figure S8.**

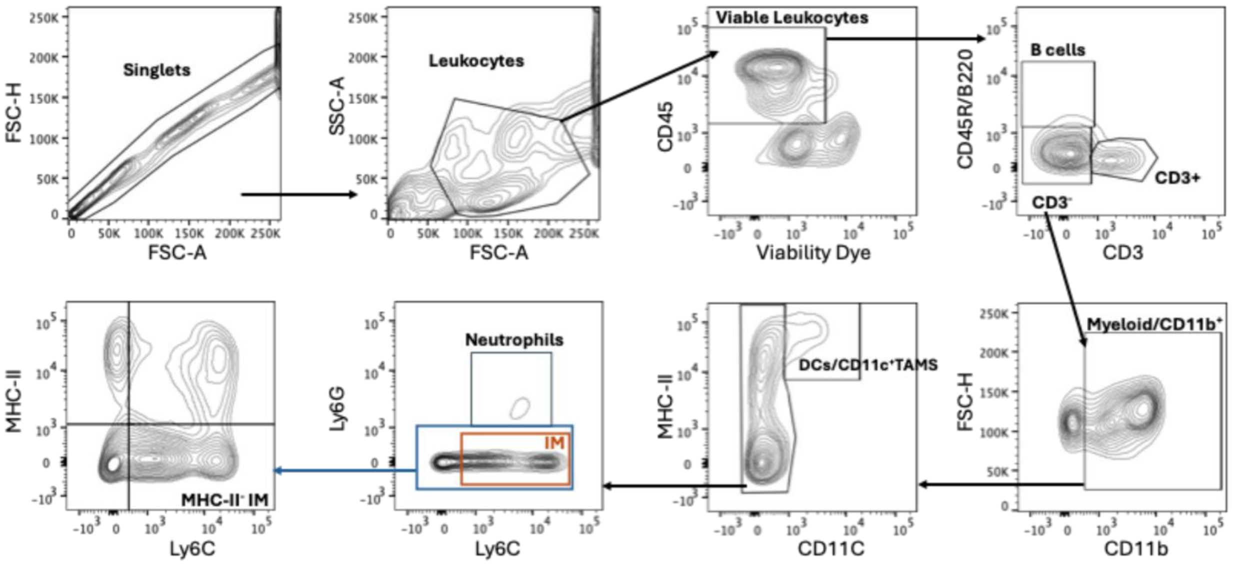


**Figure S8. Flow-cytometry gating strategy for myeloid populations infiltrating the bladder.** Representative flow-cytometry plots showing the sequential gating strategy used to characterize bladder-infiltrating myeloid populations. Singlets were selected using FSC-A and FSC-H, followed by leukocyte selection using FSC-A and SSC-A and identification of viable $CD45^+$ cells. $CD3^+$ T cells and $CD45R/B220^+$ B cells were excluded to define the CD3- compartment, and from which $CD11b^+$ myeloid cells were selected. Following exclusion of $CD11c^+$ cells, neutrophils were identified as $Ly6G^+$ cells and inflammatory monocytes (IM) as $Ly6G^-Ly6C^+$ cells. The $Ly6G^-$ compartment was further resolved according to Ly6C and MHC-II expression to distinguish $Ly6C^+MHC\text{-}II^-$/low IM.

**Figure S9.**

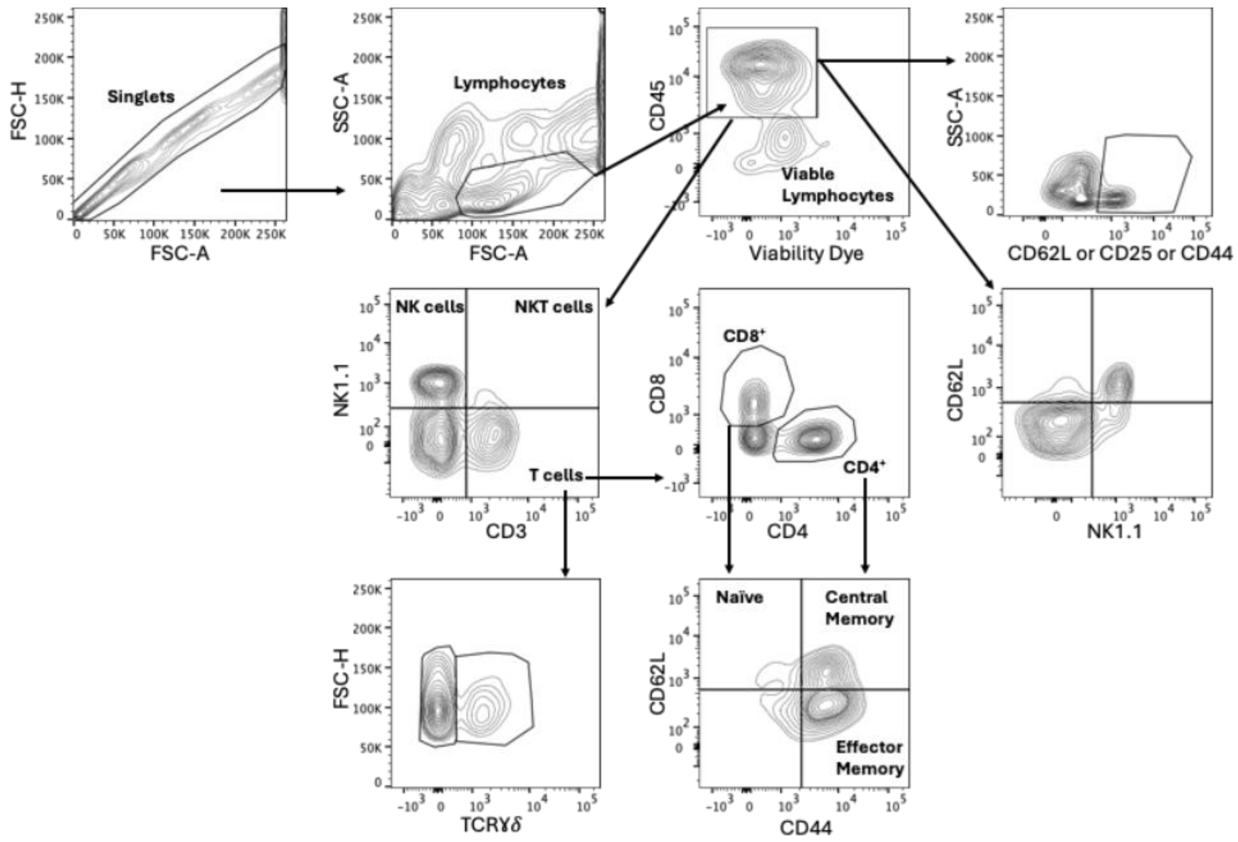


**Figure S9. Flow-cytometry gating strategy for lymphoid populations infiltrating the bladder.** Representative flow cytometry strategy used to identify leukocytes infiltrating the bladder. Singlets were first selected, followed by lymphocyte gating based on forward and side scatter parameters. Viable CD45$^+$ tumor-infiltrating leukocytes were identified using a fixable viability dye. T cells (CD3$^+$), NK cells (NK1.1$^+$CD3$^-$), NKT cells (CD3$^+$NK1.1$^+$) and TCRγδ$^+$ cells were subsequently defined. Additionally, the frequencies of CD62L$^+$, CD25$^+$, and CD44$^+$ cells were quantified within the indicated populations. CD4$^+$ and CD8$^+$ T-cell subsets were identified within CD3$^+$ cells. Memory subsets were characterized based on CD44 and CD62L expression, distinguishing naïve (CD44$^-$CD62L$^+$), central memory (CD44$^+$CD62L$^+$) and effector memory (CD44$^+$CD62L$^-$) populations.

**Table S1**. Antibody panels used in the study for cytometry studies.

| **PANEL 1** | | | |
|---|---|---|---|
| **Antibody** | **Clone** | **Fluorochrome** | **Source** |
| CD45 | 30F11 | PerCP | BioLegend |
| CD3 | 145-2c11 | APC-Cy7 | BioLegend |
| CD4 | RM4-5 | FITC | BioLegend |
| CD8 | 53-6.7 | Alexa 700 | BioLegend |
| CD62L | MEL-14 | APC | BioLegend |
| NK1.1 | PK136 | BV650 | BioLegend |
| CD127 | A7R34 | BV786 | BioLegend |
| CD44 | IM7 | PE-Dazzle | BD |
| CD25 | PC61 | PE | BioLegend |
| TCRγδ | GL3 | BV421 | BioLegend |
| Fixable Viability | | eFluor 506 | ThermoFisher |

| **PANEL 2** | | | |
|---|---|---|---|
| **Antibody** | **Clone** | **Fluorochrome** | **Source** |
| CD45 | 30F11 | PerCP | BioLegend |
| CD3 | 145-2c11 | APC-Cy7 | BioLegend |
| CD45R/B220 | RA3-6B2 | FITC | BioLegend |
| CD11b | M170 | BV650 | BioLegend |
| CD11c | HL3 | APC | BD Pharmingen |
| Ly6G | 1A8 | BV421 | BD Pharmingen |
| Ly6C | AL-21 | PE | BD |
| MHCII | M5/114.15.2 | BV605 | BioLegend |
| Fixable Viability | | eFluor 506 | ThermoFisher |
